# H3K27me3 Drives Constricted Migration–Induced 3D Genome Rewiring in Melanoma

**DOI:** 10.64898/2026.09.11.751064

**Authors:** Taiwo Habeeb Olajide, Muhammad Adeyemi Ajagbe, Alvaro Rodriguez Gonzalez, Hameed Ijadunola, Chandler Ross, Lily Eisenklam, Amanda Swets, Rachel Patton McCord

**Affiliations:** Department of Biochemistry & Cellular and Molecular Biology, University of Tennessee Knoxville, Knoxville, TN 37996, USA; Genome Science and Technology Program, Bredesen Center, University of Tennessee Knoxville, Knoxville, TN 37996, USA; Department of Electrical Engineering and Computer Science, University of Tennessee Knoxville, Knoxville, TN 37996, USA

## Abstract

Repeated exposure of neoplastic cells to mechanical stresses during metastasis can drive stable morphological and physiological changes. We previously showed that A375 human melanoma cells subjected to 10 rounds of constricted migration (“Bottom-10 cells”) became significantly more migratory and exhibited H3K9me3 relocalization, transcriptional remodeling, and chromatin compartment changes relative to naïve parental A375 cells. Because these phenotypes persist across cell divisions, we hypothesized they are driven and maintained by specific epigenetic mechanisms.

We profiled candidate histone modifications by CUT&RUN and applied a computational model to identify marks that explain compartment switches. In parental cells, H3K9me3 and H3K4me1 strongly predict locus stability in the B and A compartments, respectively. In contrast, changes in H3K27me3 dynamically predict A-to-B compartment switching after constricted migration. Consistently, altered H3K27me3 enrichment correlates with both differentially regulated genes and chromatin compartment changes between Bottom-10 and parental populations. We therefore tested whether inhibiting H3K27me3 deposition alters migratory potential and/or the “memory” of the increased migratory phenotype.

Inhibiting EZH1/2 reduced migration efficiency overall, with a stronger effect on constricted than unconstricted migration. This reduction was accompanied by decreased genomic H3K27me3 enrichment and reversal of some compartment changes in Bottom-10 cells. However, acute treatment did not durably reverse the highly migratory Bottom-10 phenotype; cells returned to their initial state after drug removal. Chronic inhibition across many sequential rounds of constricted migration dramatically reduced the fraction of cells able to migrate. Notably, cells that migrated despite chronic EZH inhibition appeared to escape treatment, acquiring many typical Bottom-10 H3K27me3 and compartment changes even though acute inhibition tends to prevent them. Overall, these results highlight H3K27me3 as a key contributor to establishing and maintaining constricted migration–induced 3D genome changes.

## Background

Cancer remains one of the leading causes of mortality worldwide, despite significant advances in our understanding of tumor biology and the development of cancer therapeutics ^1,2^. Metastasis, the defining hallmark of malignant progression, accounts for the largest proportion of cancer-associated deaths^3,4^. During metastasis, neoplastic cells must often squeeze through constrictions smaller than their nuclei to access and colonize secondary sites^4,5^.

We previously established an experimental metastasis-inspired model using A375 melanoma cells in a 2D constricted-migration assay with 5 μm pores. After ten consecutive rounds of constricted migration, the successfully migrated population (“Bottom-10” cells) displayed altered overall morphology and a stable increase in migration efficiency^6^. Prior work has shown that mechanical forces, including constricted migration, confined migration, shear stress^7^, stretching^8^, substrate stiffness^9^, and compression^10^, can remodel chromatin state and higher-order genome organization, producing changes in cell fate, heterochromatin marks, chromatin accessibility, and transcriptional programs. Some of these effects are stable, while others are transient^6,11–13^. Consistent with this literature, A375 Bottom-10 cells in our system stably exhibit not only the previously described morphological phenotypes^14^, but also transcriptional and genome conformation changes. Persistent chromatin remodeling coupled to durable phenotypic change, as observed in Bottom-10 cells, has been proposed to reflect an underlying form of epigenetic memory^15,16^. However, it remains unclear whether A375 Bottom-10 cells establish a distinct epigenetic landscape that underlies both the acquisition and maintenance of their novel morphology and 3D genome conformation.

Polycomb repressor complexes (PRCs), and in particular Polycomb repressive complex 2 (PRC2), mediate mono-, di-, and trimethylation of histone H3 lysine 27 (H3K27) through the catalytic subunits EZH1 and EZH2, generating the repressive H3K27me3 mark. H3K27me3 has been shown to co-occupy promoters of developmental and lineage-determining genes with H3K4me3, and its cell-specific activation or repression plays a significant role in developmental transitions and cell-fate determination ^17,18^. This mark is broadly implicated in gene silencing and has been associated with cancer progression and aggressive/metastatic phenotypes across multiple tumor types^19,20^. Enhancer of Zeste Homolog 2 (EZH2) is an important regulator of melanoma progression, with elevated EZH2 expression and activity associated with aggressive disease, invasion, and metastatic potential ^21–23^. Notably, H3K27me3 has also been reported to change dynamically in response to mechanical cues. H3K27me3 increased in stem cells cultured on softer substrates and decreased when cultured on stiffer substrates^9^, while its levels increased in neoplastic models under confinement^24^, suggestive of a cell type-specific mechanosensitive epigenetic response.

Several oncotherapeutics have been developed to target epigenetic regulators. Tazemetostat, an FDA-approved EZH2 inhibitor, has demonstrated antitumor activity in melanoma by inducing STING expression and thereby enhancing antitumoral immunity^25^. However, emerging evidence suggests that loss of EZH2 activity can drive compensatory increases in EZH1 translation and activity^26^. While EZH2 inhibition has been reported to reduce A375 migration through 8 μm pores^27^, it remains to be determined whether EZH inhibition is capable of reversing the high migration efficiency and 3D genome topology in A375 Bottom-10 cells, which acquire a stable, highly migratory phenotype after ten rounds of constricted migration.

In this study, we sought to determine whether highly migratory A375 Bottom-10 cells exhibit an altered epigenetic landscape relative to Parental cells, and to define how any such differences contribute to the acquisition and maintenance of the Bottom-10 migratory phenotype and associated chromatin remodeling. We showed that joint inhibition of EZH1 and EZH2 reduced the migratory efficiency of melanoma cells, partially reversed chromosome compartment changes induced by sequential rounds of constricted migration, and hindered the complete acquisition of the epigenetic and 3D genome features of highly migratory melanoma cells.

## RESULTS

### Pre-existing histone modification patterns in Parental A375 cells accurately predict chromosome compartment changes in Bottom-10 cells

Studies have established the alignment of histone epigenetic marks with genome compartment identity^28^ and demonstrated how using histone enrichment data in a supervised learning model can accurately predict genome compartment identities^29^. We have previously reported differential 3D genome topology between parental and highly migratory Bottom-10 A375 cells, demonstrating stable chromatin remodeling^6^. Here, we asked whether histone modifications can be used to explain these chromosome compartment changes. Our previous work noted that there was no long-term change in H3K9me3 intensity in Bottom-10 cells^6^ but other studies have reported an increase in H3K27me3 immediately after constricted migration^24^. We checked H3K27me3 levels immediately after A375 cells had passed through 5 rounds of constricted migration, at which point the migratory phenotype is similar to Bottom-10 cells. In comparison to the unmigrated cells, we observed a slight increase in the intensity of H3K9me3 and H3K27me3 and a change in the distribution of the signal in our migrated cells (Supp. Fig.1 a-c).

**Fig 1:**
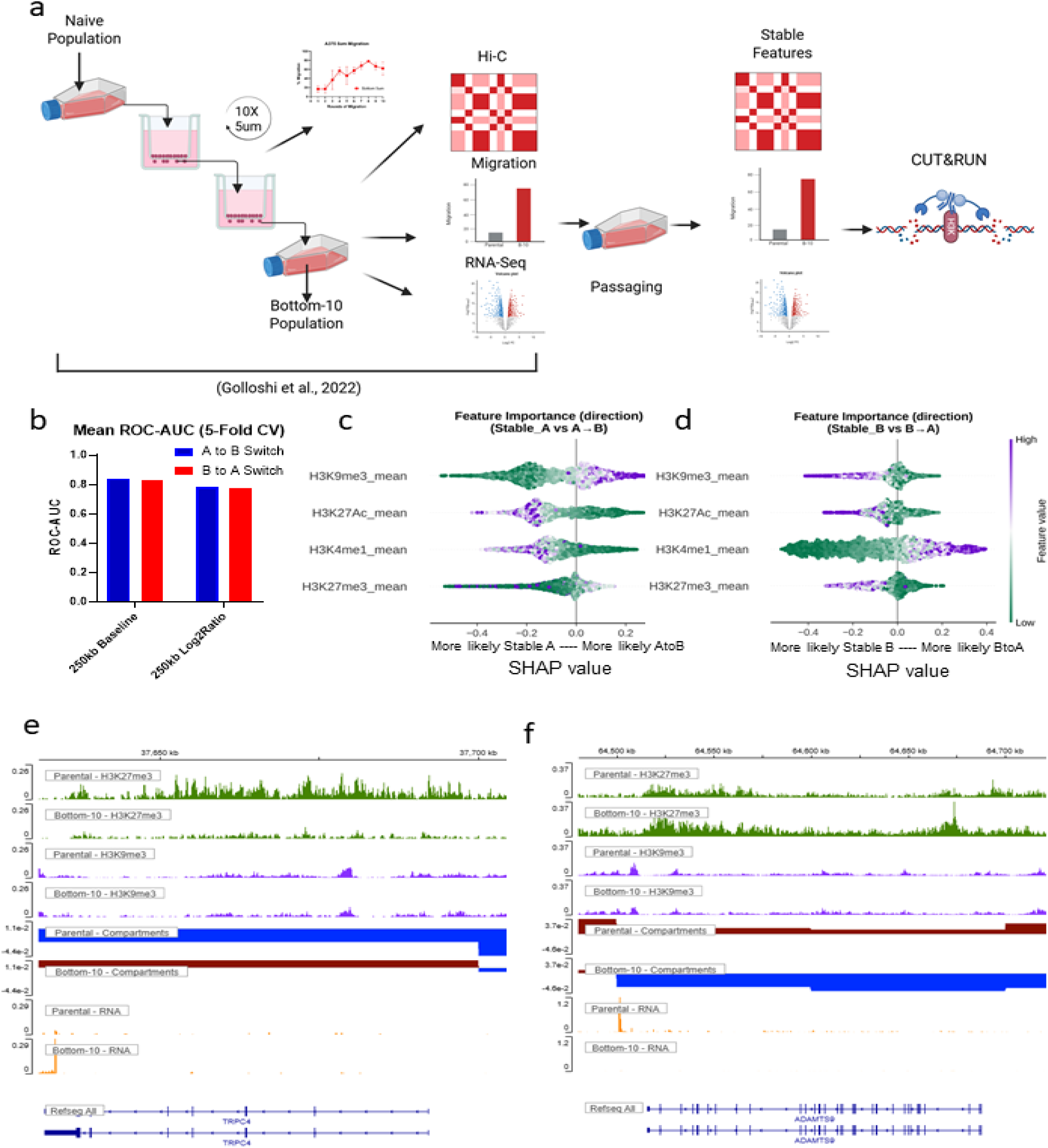
Histone modification patterns in melanoma cells predict locations of compartment switches after ten rounds of constricted migration. **(A)** A375 Melanoma cells subjected to ten rounds of sequential migration through 5 μm 2D membrane pores exhibited stable differences in 3D genome, transcriptomic, and migration profiles in comparison to the naïve unmigrated cells. We surveyed four histone epigenetic markers between the Migrated Bottom-10 cells and the unmigrated parental cells. **(B)** Bar plot showing the accuracy of prediction of the chromatin compartment changes in the Bottom-10 vs. Parental cells based on combining enrichment patterns of four histone epigenetic markers in a Random Forest model. Beeswarm SHAP plots show locus-level feature contributions of each histone mark to the predicted probability of compartment switching from Parental to Bottom-10, either A to B **(C)** or B to A **(D)**. Positive values indicate that the given level of that histone mark increases the probability of the switch, whereas negative values support the corresponding stable-compartment class. Features are ranked by mean absolute SHAP value within each model. Point color denotes the feature value (signal of the given histone mark in Parental cells in each 250 kb bin) from low (green) to high (yellow) **(E)** Genome track showing the Parental vs. Bottom-10 changes in the enrichment pattern of H3K27me3, H3K9me3, Compartment Identities, and gene transcription for the upregulated gene TRPC4. A decrease in H3K27me3 corresponds to a switch from B (blue) to A (red) and an increase in RNA signal at this gene, while H3K9me3 remains unchanged. **(F)** Same as (E), but showing an increase in H3K27me3 associated with an A-to-B switch and a decrease in RNA transcription at the gene ADAMTS9.

However, changes in these visible patterns cannot reveal which regions experience increases in these modifications or whether there is a redistribution of the mark from some regions to others. Therefore, using CUT&RUN, we assayed the enrichment of H3K9me3, H3K27me3, H3K4me1, and H3K27ac in the Parental and Bottom-10 populations to compare with our previously collected Hi-C compartment profiles and gene expression changes (Fig. 1a). To test if the initial epigenetic state can predict the chromatin remodeling that we see in Bottom-10 cells, we trained Random Forest models on the mean signal of these marks across 250kb bins in the Parental cells to predict compartment changes in the Bottom-10 cells. Our models attained an accuracy of 84% (AUC=0.839 +/− 0.015) in predicting bins that switched compartments from A to B and 78% (AUC=0.775 +/− 0.012) for B to A (Fig. 1b and Supp. Fig. 1d). Using the change in chromatin state between Parental and Bottom-10 cells was also effective at predicting compartment changes (Fig.1b, “250kb Log2Ratio”).

To ensure that our models learned from the epigenomic marks rather than only inferring the initial spatial location of genomic loci, we tested the prediction of the models on marks only, PC1 values only, and both together. The model performed better on marks only, independent of locus location, on bins switching from B to A (marks only=78% and PC1 only=64%) (Supp. Fig.1 e). The initial compartment state (PC1 only) was a strong predictor of which bins would switch from A to B, but adding histone mark data further improved prediction accuracy. Overall, this analysis shows that there are epigenetic states in the Parental cells that predispose regions to switch during constricted migration.

To determine which epigenetic marks most strongly predict compartment stability versus switching, we performed SHAP analysis on the classifiers for each transition direction (Fig. 1c–d). Loci that are stable in the A compartment have higher enrichment of H3K4me1 and H3K27ac, while high H3K9me3 in an A compartment region increases the chance that the region will shift to the B compartment after constricted migration. H3K27me3 had a bimodal effect, where the bins with the highest enrichment were likely to remain in the A compartment while moderate levels of H3K27me3 predicted shifts toward the B compartment. (Fig. 1c). Loci stable in the B compartment tended to have high enrichment of H3K9me3 and low enrichment of H3K4me1, while high levels of H3K4me1 enrichment predicted switching from the B to A compartment after constricted migration. (Fig. 1d). Overall, we see that if a genomic region in Parental cells is marked with histone modifications that conflict with its spatial compartment (heterochromatin marks in the A compartment or euchromatin marks in the B compartment), these regions are primed to switch compartments during constricted migration, bringing their histone profile into concordance with their compartment association.

### H3K27me3 enrichment levels correlate highly with transcriptional changes and compartment switches in A375 melanoma cells

Three-dimensional genome topology is an additional layer of transcriptional regulation that is closely coupled to epigenetic chromatin states; its spatial organization can influence gene expression through the distribution and activity of histone modifications and other epigenetic features^30^. Along with 3D structural changes reported in Bottom-10 cells, we have also reported transcriptional changes. Therefore, we asked whether histone mark changes align with both genome compartment changes and transcriptional changes in our system. Taking the example of TRPC4, a gene that was significantly upregulated in Bottom-10 cells compared with parental cells, we saw no difference in the comparative enrichment of H3K9me3, but there was a marked reduction in the enrichment of H3K27me3 across the gene body in Bottom-10 cells. This change coincides with the A-to-B compartment switch from Parental to Bottom-10 cells (Fig. 1e). Across the gene body of ADAMTS9, a downregulated gene in Bottom-10 cells, there was an increase in the enrichment of H3K27me3 in Bottom-10 cells compared with parental cells. This change aligns with a genome compartment switch from A to B in Bottom-10 cells (Fig. 1f).

To determine whether these locus-specific observations reflect a broader trend, we extended the analysis to the full set of differentially expressed genes and genomic loci that switch compartments in Bottom-10 cells; we quantified how histone enrichment differs between Parental and Bottom-10 populations. Using the ComputeMatrix function on the Galaxy platform^31,32^, we observed that H3K27me3 changes were most consistently associated with both transcription and compartment changes in both activating and repressing directions (Figure 2 and Supp. Fig 2). Relative to Parental cells, genes downregulated in Bottom-10 cells and regions that switch from A to B show higher average H3K27me3 enrichment in Bottom-10 cells (Fig. 2a and c). Conversely, genes upregulated in Bottom-10 cells and regions that switch from B to A show lower average H3K27me3 enrichment (Fig. 2b and d). In contrast, H3K9me3 patterns change very little on average for these regions of differential expression or compartment switches (Fig. 2e-h). Active chromatin-associated marks H3K4me1 and H3K27Ac also change in directions that correlate with changes in gene expression, increasing around upregulated genes and decreasing around downregulated genes (Supp. Fig 2). But changes in these activating marks are less strongly associated with compartment changes, in particular showing no significant changes in regions with B-to-A compartment shifts (Supp. Fig. 2d and h).

**Fig 2:**
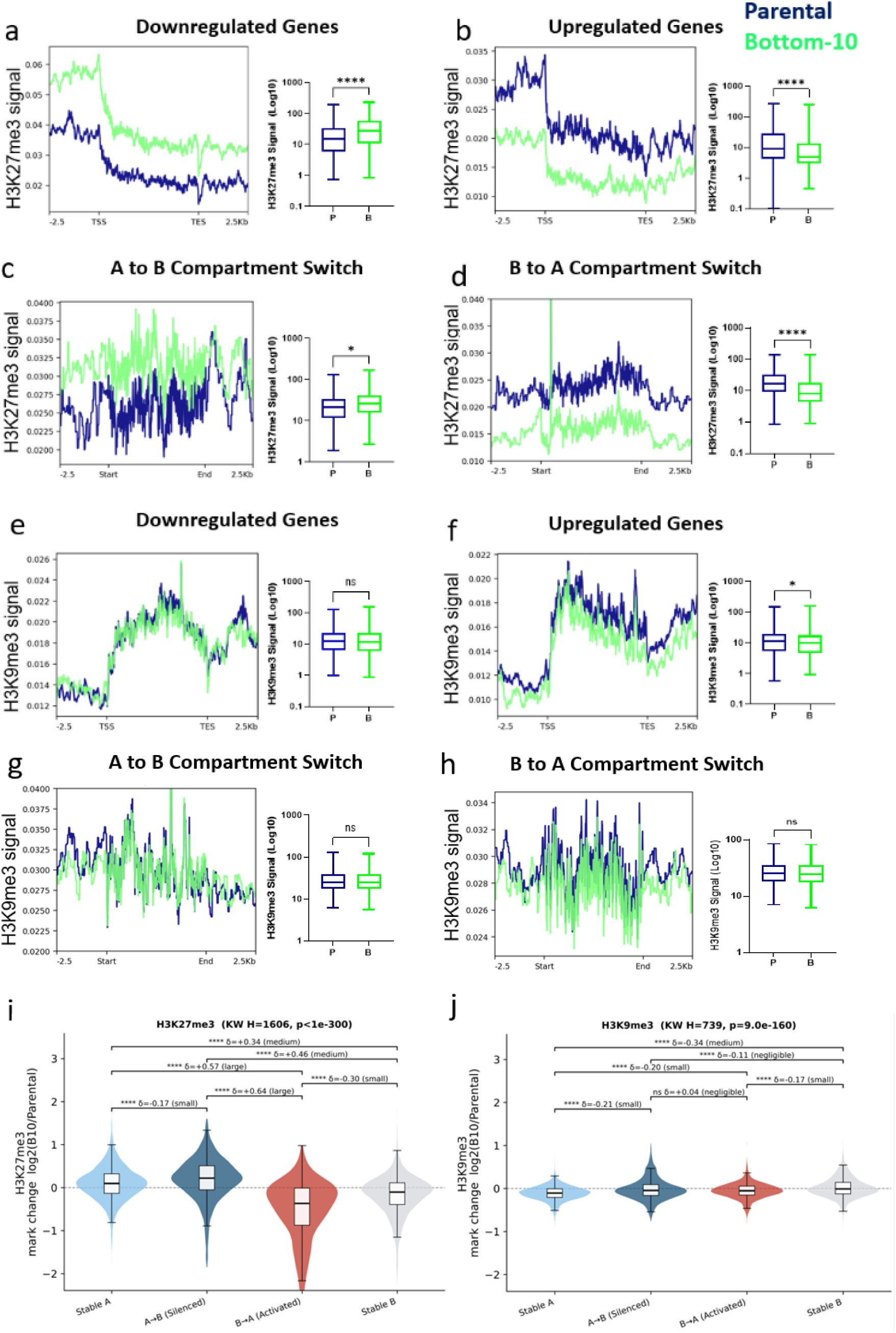
H3K27me3 enrichment levels strongly correlate with transcriptional changes and compartment switches in A375 melanoma cells. (a-h) Pile-up plots show the average histone modification signal in Parental (blue) or Bottom-10 (green) cells across sets of differentially regulated genes scaled from transcription start to transcription end (a, b, e, f) or across sets of 250 kb bins that switch compartments (c, d, g, h). H3K27me3 signals are shown in a-d while H3K9me3 signals are shown in e-h. Paired boxplots show the distribution of signals across the sets of regions averaged in the pileup. Each point that contributes to the boxplot is the average signal across one gene or one 250 kb bin. Boxplots show the minimum-to-maximum range, quartiles, and median. Significance is calculated by Welch’s t-test (* p<0.05, ** p<0.01, *** p<0.001, **** p<0.0001) (i) Violin plot showing log2 fold change in H3K27me3 signal from Parental to Bottom-10 cells across 250 kb bins categorized according to whether they show stable or altered compartment state between Parental and Bottom-10 cells. The significance (stars as described above) is calculated using the Kruskal–Wallis test, and the effect size is calculated using Cliff’s delta. (j) same as (i) but for H3K9me3 signals along the genome.

Next, we determined the effect size of these marks’ contribution to chromatin compartment switches and compared stable and switching loci. To do this, we extracted mean enrichment for each mark in 250 kb bins and computed the change between Bottom-10 and Parental cells as log_2_(B10/Parental). Our analysis showed that H3K27me3 strongly separated the four bin classes (Kruskal–Wallis, p < 1× 10^−300^) and exhibited similar associations with locus identities as previously described. When we compared the bins that switched to A against those that shifted to B, we see a large effect size difference (δ = +0.64, large, p = 4.4 × 10⁻¹⁸⁵). When the gain of H3K27me3 in A→B bins and loss in B→A was compared to Stable B bins, the effect size differences of δ = +0.46 (medium) and δ = −0.30(small), respectively, persisted, indicating that these shifts are not solely explained by the final compartment state. Both switching groups also exhibited broader distributions than the stable groups, suggesting substantial bin-to-bin variability in H3K27me3 gain or loss (Fig. 2i). In contrast to H3K27me3, H3K9me3 showed a mirror-image pattern. (Fig. 2j). This pattern indicates that H3K9me3 tracks locus stability rather than compartment dynamics (Fig. 2j). H3K4me1 also separated the four groups strongly (Kruskal–Wallis H = 966, p = 3.5 × 10⁻²⁰⁹), but asymmetrically (Supp. Fig.2i), while H3K27ac produced the largest omnibus statistic among the four marks (Kruskal–Wallis H = 2081, p < 1 × 10⁻³⁰⁰), but with a different separation structure. The overall range of H3K27ac change was also narrower than for H3K27me3, with most bins falling within ±0.3 (Supp. Fig. 2j). Notably, H3K27me3 was the only mark for which the two switching groups were more strongly separated from each other than from any other group; for all other marks, the largest separation involved a stable group.

Taken together, evidence from these markers and their relationship to transcriptomic and chromatin compartment dynamics suggested H3K27me3 as a key marker modulating transcriptomic and 3D genome structure differences between the Parental and Bottom-10 A375 populations.

### Dual inhibition of EZH1 and EZH2 reduces constricted migration efficiency in A375 melanoma

We next decided to pharmacologically inhibit the deposition of H3K27me3 in A375 cells to assess whether this reverses the differential phenotypic characteristics associated with Bottom-10 cells. Although both catalytic subunits of the PRC2 are involved in the methylation of the lysine tail of H3K27, evidence has suggested compensatory activity of EZH1 when EZH2 is knocked down^26^. Thus, we tested joint inhibition of EZH1 and 2, as well as EZH2-specific inhibition (Fig. 3a). To mimic transwell experimental conditions, we assessed the viability of A375 cells under different dosages by treating the cells in fully supplemented media for 24 hours, followed by treating for another 24hours in unsupplemented media before conducting an MTS assay. In comparison to the vehicle treatment, UNC1999 treatment marginally increased the metabolic activity of the cells up to 5μM concentration, while activity was comparable 10μM. EPZ005687 treatment doses didn’t result in any change in the metabolic activity of the cells (Supp. Fig.3 a-b). After establishing the viability of the cells under differing dosages of the agents, we assessed the cell response to the inhibition. As seen in the representative image and the quantification boxplots, it was determined that there was dose-dependent depletion of H3K27me3 levels in both parental and Bottom-10 cell populations when treated with EPZ005687 (Supp. Fig. 3c-d). In response to the joint inhibition of EZH1 and EZH2 with UNC1999, there was a reduction in the fluorescence intensity of H3K27me3 in both A375 populations (Fig. 3b). This reduction reached statistical significance at 5 μM in both parental and Bottom-10 cells and hence this inhibitor concentration was used for the remaining experiments (Fig. 3c).

**Fig. 3:**
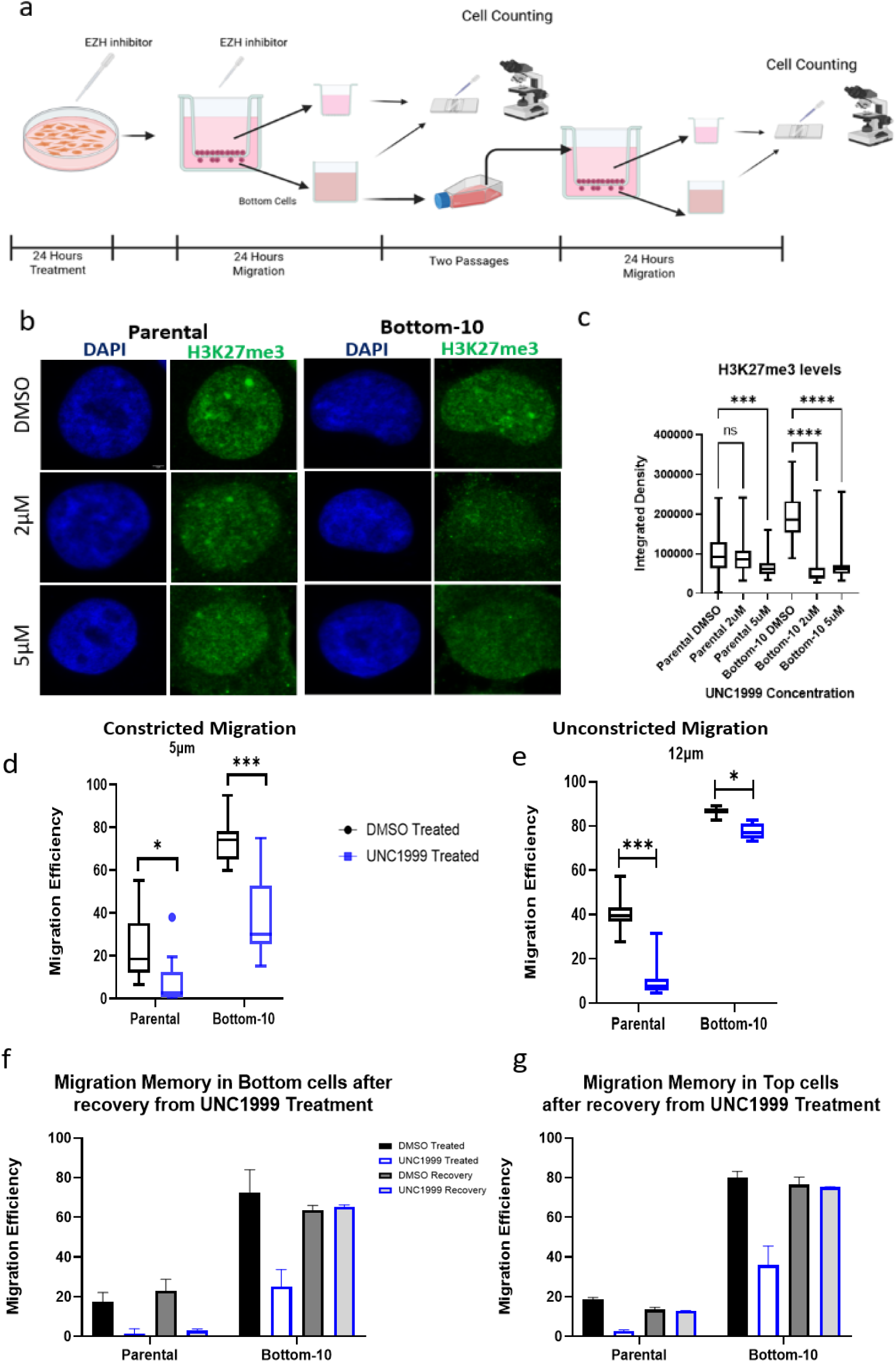
Inhibition of EZH1 and EZH2 reduces migration efficiency in A375 melanoma cells. (A) Schematic of experiment: A375 cells were treated with 5 μM UNC1999 in fully supplemented media for 24 hours, followed by another 24 hours of treatment while undergoing constricted migration. The collected cells were allowed to recover for two passages before being subjected to constricted migration to assess migration memory. (B) Representative fluorescent image showing Parental and Bottom-10 cells treated with increasing doses of UNC1999. (C) Quantification plots show the integrated density of H3K27me3 signals in both the parental and Bottom-10 A375 cell populations after treatment with UNC1999. (D) Box plot showing the migration efficiency of A375 parental and Bottom-10 cells treated with UNC1999 migrating through a constricted migration (5 μm pores). Each box represents at least 12 replicates. P values refer to t-tests. (* p<0.05, *** p<0.001) (E) Same as D but for unconstricted migration (12 μm pores). Each box represents 9 replicates. (F) Bar plot showing the migration efficiency of cells during UNC1999 treatment and then after recovery for the cells that migrated (“Bottom” of the Transwell) during that treatment. N = 3 replicates. (G) Same as F for a replicate experiment of UNC1999 migration, then testing the migration of the cells that did not migrate (“Top” pf the Transwell) during the initial treatment test. N=3 replicates.

Next, we evaluated the effect of UNC1999 treatment on constricted migration efficiency in both A375 populations. In both naïve parental and Bottom-10 A375 cells, there was a significant reduction in migration efficiency after the 24-hour EZH inhibition. Notably, in some replicates of the naïve parental cells, there was complete inhibition of migration (Fig. 3d). EZH2-specific inhibition, however, didn’t significantly reduce the migration efficiency in either A375 population (Supp. Fig. 3e). We further assessed whether the reduction in migration in response to UNC1999 inhibition is constriction-dependent. Using the same experimental setup with 12µm pores in lieu of the 5 µm pores, we recorded a reduction in migration efficiency in both cell populations (Fig.3 e). However, the magnitude of the reduction is much lower in the Bottom-10 cells, suggesting that EZH inhibition specifically affects the ability of these highly migratory cells to migrate through nucleus-constricting pores.

We previously showed that the Bottom-10 cellular, nuclear, and migratory phenotypes that arise after constricted migration are stable. So, we next checked whether acute inhibition of EZH1 and EZH2 eliminates migration stability in these cells. The Bottom-10 cells collected from the EZH1&2i-constricted migration setup were cultured through two passages to recover from the inhibition treatment and challenged through a subsequent Transwell constricted migration without treatment. The parental cells that passed through constrictions under EZH inhibition treatment remained very poor at migrating even after a treatment recovery period (Fig. 3f). This long term migration reduction was not seen for the parental cells that were treated but did not migrate during treatment (Fig. 3g). This suggests that the combined effect of constricted migration stress during EZH inhibition in the parental cells caused a long term reduction in migratory capacity for these cells. However, in Bottom-10 cells, for both migrated and unmigrated populations, migration capacity returns to a high level after recovery from EZH inhibitor treatment (Fig 3f and g). Therefore, while the acute EZH inhibitor treatment influences migration, it does not reprogram these cells or cause them to “forget” their highly migratory phenotype.

Taken together, we see that joint inhibition of EZH1 and 2 reduces migration in A375 cells, and the inhibition-induced reduction in migration efficiency is constriction-specific. Acute EZH1 and 2 inhibition is insufficient to deprogram the migration memory in Bottom-10 cells.

### Acute EZH1 & 2 inhibition causes significant epigenomic and transcriptional changes in A375 melanoma cells

Co-inhibition of EZH1 and EZH2 resulted in a pronounced effect on the migratory phenotype of A375 cells, so we next asked what effect this acute treatment had on the H3K27me3 mark distribution, gene expression, and genome folding patterns in these cells, particularly those which are known to shift with constricted migration. Using CUT&RUN, we measured the effect of UNC1999 on the genomic enrichment of H3K27me3 at the previously identified differentially expressed genes (DEGs) in Bottom-10 cells compared to Parental cells and at loci that switched compartments in the Bottom-10 population vs. Parental. Along the gene bodies of DEGs, we observed a reduction in H3K27me3 in the UNC1999-treated conditions for both the Parental (PU) and Bottom-10 populations (BU). This reduction in H3K27me3 after treatment is much more significant in regions that shift toward the B compartment or are downregulated after constricted migration than in regions that increase transcription or shift to the A compartment (Fig. 4a-d). This suggests that the decrease in H3K27me3 that results from EZH inhibition particularly impacts regions which increase their H3K27me3 and move toward a repressive B compartment state after constricted migration.

**Fig 4:**
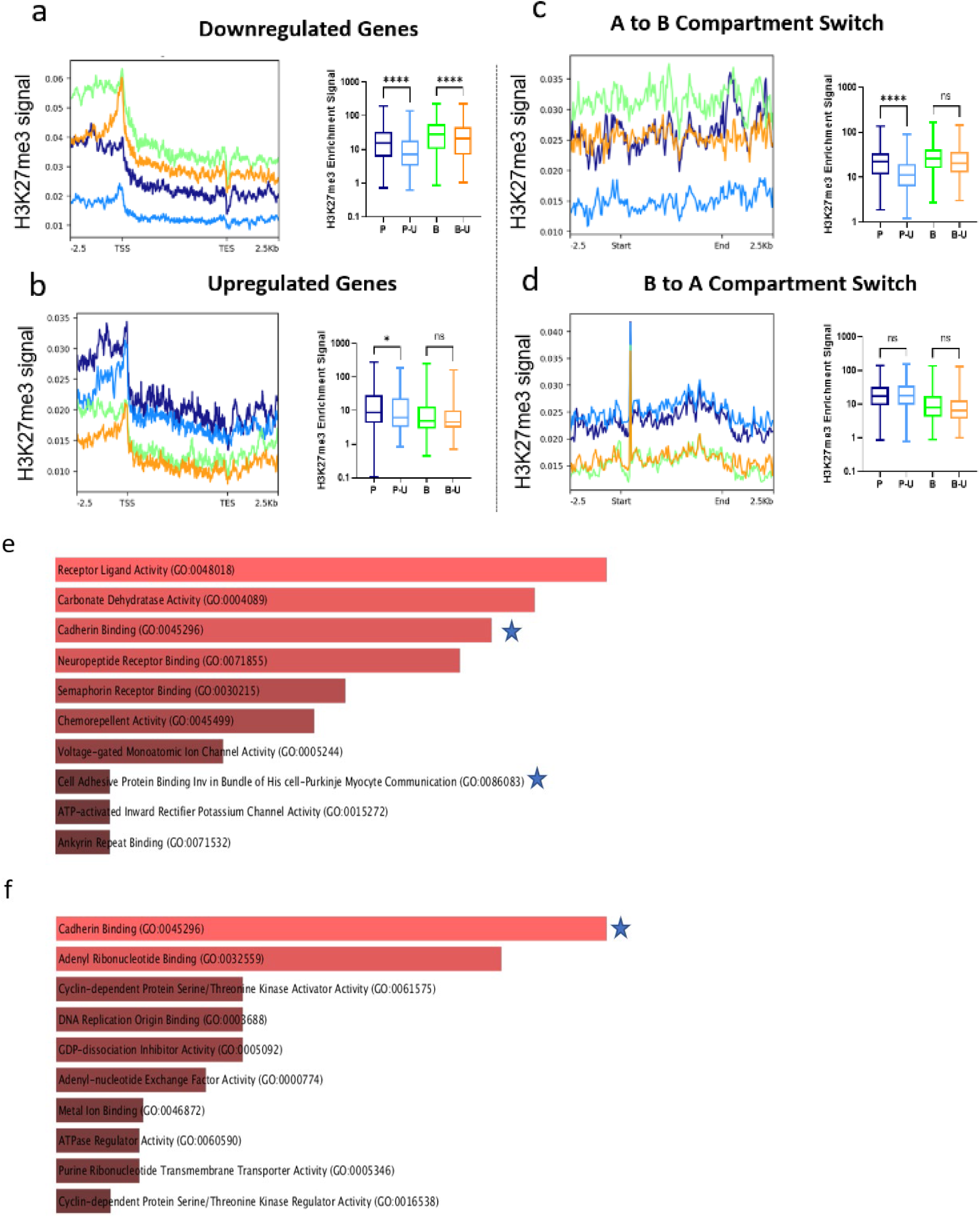
Acute EZH1 & 2 inhibition led to significant epigenomic and transcriptional changes in A375 Melanoma cells. Average line plots and box plots showing the CUT&RUN signal for H3K27me3 with and without UNC1999 treatment across the gene body of genes previously identified to be downregulated (A) or upregulated (B) after 10 rounds of constricted migration. P=Parental, PU=Parental treated with UNC1999, B = Bottom-10, BU = Bottom-10 treated with UNC1999. (C and D) Same as A and B, but for 250 kb loci previously identified to switching from the A to B compartment (C) or B to A compartment (D) in the A375 Bottom-10 cells compared to Parental. Significance is calculated by Welch’s t-test (* p<0.05, ** p<0.01, *** p<0.001, **** p<0.0001) (E and F) Gene Ontology Molecular Function enrichment (Calculated by EnrichR) for the set of genes measured by RNA-seq to be upregulated in Parental (E) or Bottom-10 (F) cells after 48 h treatment with UNC1999. (F) Gene Ontology of the upregulated genes in the Bottom-10 A375 cells after acute UNC1999 Treatment. Bar length indicates enrichment score while color saturation indicates significance. Terms of interest are starred.

Next, we asked if the change in the enrichment of H3K27me3 translated to an altered transcriptional state of the cells, which might help explain a change in migratory phenotype. Comparing RNA-seq data between vehicle and UNC-treated A375 parental cells revealed upregulated genes whose gene ontology analysis identified processes such as Cadherin Binding and Cell Adhesive Binding (Fig. 4e). The comparison between treated and untreated Bottom-10 cells also revealed upregulated genes that are involved in molecular processes such as Cadherin binding as well (Fig. 4f). An increase in these genes could promote cell adhesions that counteract migration. Genes involved in cellular immune response populated the downregulated genes in both cell populations. (Supp. Fig. 4 a-b)

### Acute EZH1 & 2 inhibition led to a reversal in some compartment changes seen between naïve parental and sequentially constricted A375 Melanoma cells

After establishing the effect of EZH inhibition on the morphological, transcriptomic, and epigenomic landscape of A375 cells, considering the difference in 3D genome structure between naïve parental and experienced Bottom-10 A375 populations, we probed effects of EZH inhibition on the 3D genome topology of A375 cells. As previously established by Golloshi et al. (2022), we performed Hi-C on treated and untreated cell populations, called compartments, and performed pairwise comparisons between the groups (see Data S1 for all quantitative compartment classifications and comparison categories).

In the Parental population, among regions that experienced compartment changes, UNC1999 resulted in a higher number of compartments drifting toward the B compartment, with a larger proportion experiencing a 10% (weak) shift in the B direction without actually changing compartment identity. In contrast, among the Bottom-10 populations, UNC1999 resulted in a larger fraction drifting towards the A compartment, with 159 loci switching from the B to the A compartment (Figure 5 a and b). Overall, UNC1999 treatment resulted in fewer compartment changes within individual populations in comparison to the pre-existing differences between Bottom-10 and Parental cell populations, indicating that UNC1999 has a modest effect in reversing the 3D genome change reported in Bottom-10 cells.

**Fig 5:**
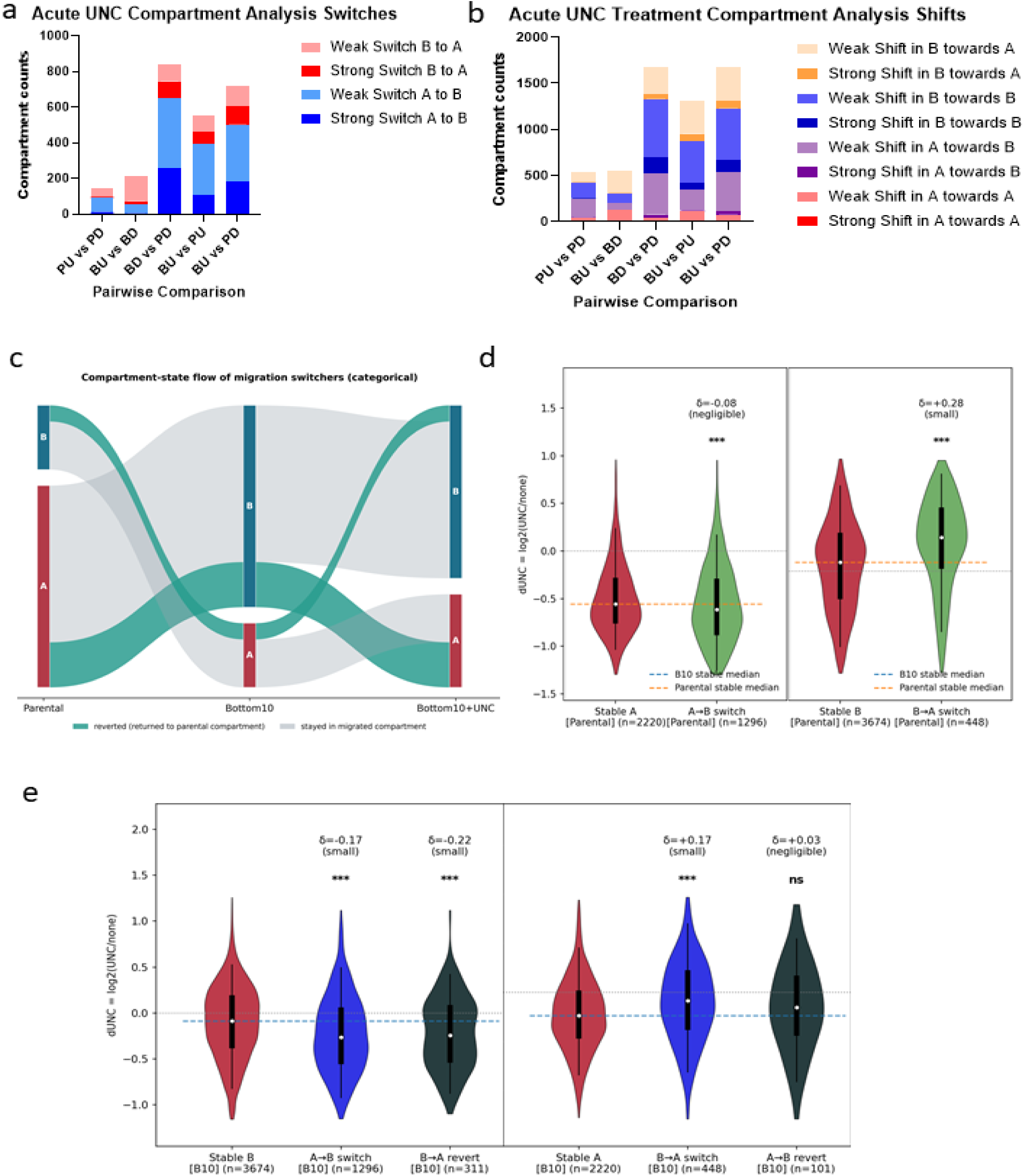
Acute EZH1 & 2 inhibition led to a reversal in some chromosome compartment changes seen between naïve parental and Bottom-10 A375 melanoma cells. (A) Stacked bar plot showing the pairwise comparison of compartment-switch categories between untreated and acute UNC1999-treated A375 cell populations. (B) Stacked bar plot showing the pairwise comparison of compartment-shift categories between untreated and acute UNC1999-treated A375 cell populations. (C) Sankey flow diagram showing genome compartment dynamics in the A375 Melanoma population in response to acute UNC1999 treatment. (D) Violin plot showing the relative effect size of Acute UNC1999 treatment on compartments in parental melanoma cells. (E) Violin plot showing the relative effect size of Acute UNC1999 treatment on compartments in Bottom-10 melanoma cells.

Furthermore, when we compared vehicle-treated and untreated conditions in both populations, we observed a significantly higher number of loci drifting toward the B compartment, consistent with earlier reports suggesting constricted migration-induced heterochromatinization. Vehicle-treated populations showed a significantly higher number of loci drifting toward the B compartment (76%), with just more than 600 loci shifting toward the B compartment by up to 10%. However, the comparison of UNC1999 treatment across both populations showed 25% fewer changing loci. Overall, treated cells showed a higher proportion of loci drifting toward the A compartment in Bottom-10 UNC1999-treated cells. To determine if UNC1999 treatment drives 3D genome conformation similarity between Bottom-10 cells and initial naïve parental cells, we compared their compartment identities, and we see that although the treatment didn’t return the genome structure of Bottom-10 to that of the parental cells, it did result in fewer overall changes in the genome structure in comparison to the vehicle-treated population. (Fig. 5a-b).

Next, we wanted to assess if UNC1999 treatment impacted the compartment identity of the previously identified loci that changed between parental and Bottom-10 cells. Tracking from naïve parental cells to Bottom-10 cells and then to UNC1999 treated Bottom-10 cells revealed that of the 1701 loci that switched to the B compartment in the Bottom-10 cells relative to the Parentals, 380 shifted back towards A after treatment with UNC1999, while of the 541 that switched from B to A in the Bottom-10 cells, 136 switched back to A after the treatment (Fig. 5c).

We then wanted to determine the concordance between the change in H3K27me3 enrichment in both populations with the compartment changes in the cells. Our analysis showed that in Parental cells, regions that will later switch during migration already exhibit similarly small, direction-dependent differences, consistent with pre-existing regional properties rather than migration-acquired targeting. Future B→A regions (B in Parental) lose less H3K27me3 than stable B regions (δ=+0.28), while future A→B regions (A in Parental) do not differ meaningfully from stable A regions (δ=-0.08). Across all comparisons, effect sizes remain small (no δ>0.28, most <0.20), and widespread statistical significance (typically p<0.001) is consistent with large bin counts (n=101–3674) rather than large biological effects (Fig. 5d).

### Acute EZH1 & 2 Inhibition led to a more significant genomic effect that explains migration inhibition in Melanoma cells

After demonstrating that EZH1&2 inhibition led to epigenomic and 3D genome structural changes and a significant reduction in the migratory capacity of A375 cells, we next assayed the genomic enrichment of H3K27me3 in cells that completed migration in both populations under the pressure of UNC1999 treatment. We compared H3K27me3 enrichment across the previously identified DEGs in Bottom-10 vs. Parental cells and for loci that switched compartments in the Bottom-10 population compared to Parental, then across the entirety of the genome. Among the Bottom-10 cell population, UNC1999 treatment during migration resulted in a significant reduction in H3K27me3 enrichment among the downregulated genes; although not significant, these genes had higher enrichment of H3K27me3 in the parental population (Fig. 6a).

**Fig 6:**
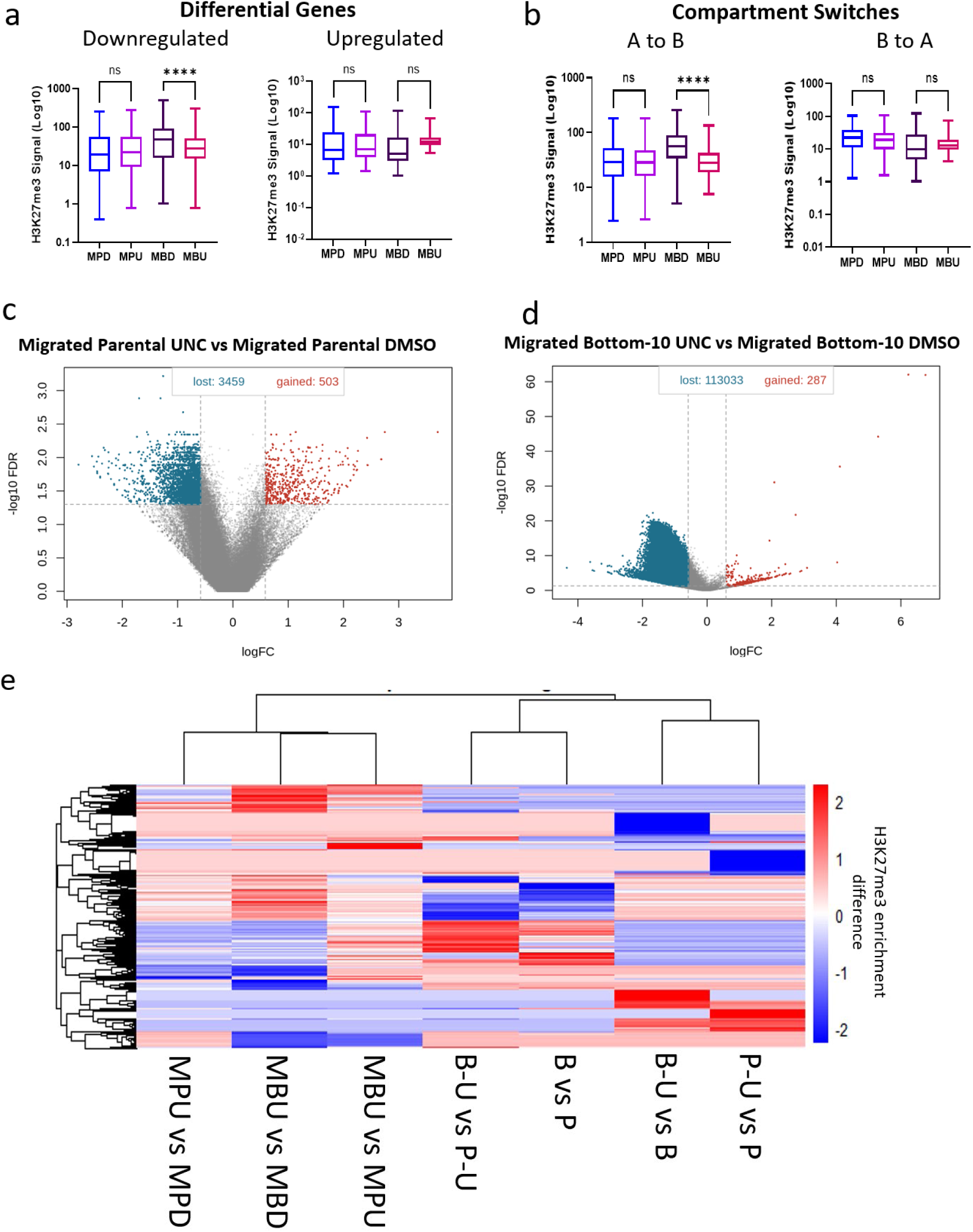
Effect of acute EZH1 & 2 inhibition combined with migration. (A and B) Box plot showing the H3K27me3 enrichment differences between migrated untreated and migrated UNC1999-treated cells in the previously identified (A) differentially regulated genes in Bottom-10 cells and (B) genome compartment switches. Values on the box plot represent bins along the whole length of the gene or compartment. (C and D) Volcano plot showing loci of differential enrichment of H3K27me3 between (C) Migrated untreated and UNC1999-treated parental cells and (D) Migrated untreated and UNC1999-treated Bottom-10 cells. (E) Heatmap showing the H3K27me3 difference at the top 1000 changing loci in pairwise comparisons of H3K27me3 enrichment in different cell populations and treatment. (P: Parentals, P-U: Parental UNC1999-treated, B: Bottom-10, B-U: Bottom-10 UNC1999-treated, MPD: Parentals Migrated + DMSO treatment, MPU: Parentals Migrated + UNC1999 treatment, MBD: Bottom-10 Migrated + DMSO Treatment, MBU: Bottom-10 Migrated + UNC1999 Treatment)

In both populations, there was a non-significant increase in H3K27me3 enrichment among the upregulated genes (Fig. 6a). In congruence with what was observed in the downregulated genes, among loci changing from euchromatin to heterochromatin, there was a marked decrease in H3K27me3 enrichment in treated and migrated Bottom-10 cells compared with untreated cells. There were no differences in H3K27me3 levels between the treated and untreated migrated parental populations (Fig. 6b). There were no differences in H3K27me3 enrichment among the comparison populations at loci switching from heterochromatin to euchromatin (Fig. 6b).

Differential analysis of H3K27me3 across the genome showed that, in comparison to the vehicle-treated cells, treated and migrated parental cells experienced reduced H3K27me3 at 3,459 loci, while treated and migrated Bottom-20 cells had reduced H3K27me3 enrichment at 113,033 loci across the genome. In contrast, treated and migrated parental cells gained H3K27me3 at 503 loci, and Bottom-10 at 287 loci (Fig. 6c-d).

Next, using a custom script (see Methods), we computed H3K27me3 locus enrichment differences between different parental and Bottom-10 treated and untreated conditions and created a matrix of the top 1000 differential loci from pairwise differences to uncover whether there is a pattern across the different pairwise comparisons. When we clustered conditions by these top differential loci, the clades are segregated by three factors: migration, UNC1999 treatment, and combined UNC1999 treatment and Migration (Fig. 6e).

Overall, we see a combined effect of migration and EZH inhibition on H3K27me3. Further, under the additional physical and selection stress of constricted migration, H3K27me3 changes are more dramatic in Bottom-10 cells compared to Parental cells.

### Chronic EZH1 & 2 inhibition across sequential rounds of migration led to the selection of a population of A375 melanoma cells that escape the effects of the drug on H3K27me3

Transwell sequential migration generates highly migratory Bottom-10 cells after ten successive rounds of constricted migration. We have previously reported that cells in our model continuously improved at migration over ten rounds, suggesting that several rounds of constriction stress are needed to encode migration memory. We therefore hypothesized that treating cells with UNC1999 over successive rounds of migration would block the steady increase in migration efficiency and the encoding of migration memory. To investigate whether EZH1 and EZH2 inhibition would prevent A375 cells from acquiring increased migratory capabilities over sequential rounds of constricted migration, we subjected the cells to UNC1999 treatment during sequential rounds of migration.

Based on the migration efficiency we recorded with acute UNC1999 treatment, which showed that parental cells migrate poorly (averaging about 9%), we adopted a 10 cm transwell migration system that could accommodate up to 15 million cells (see Methods) and therefore recover enough cells to proceed to a subsequent migration round. Cells migrated under UNC1999 treatment initially improved, but after the fourth round they progressively worsened with each successive round. Notably, cells under UNC1999 treatment migrated consistently less than cells administered with the vehicle, which improved over the rounds of migration (Fig. 7a). The sequential migration experiment was halted at round 8, as the count of migrated cells was too low to continue with the migration protocol.

**Fig 7:**
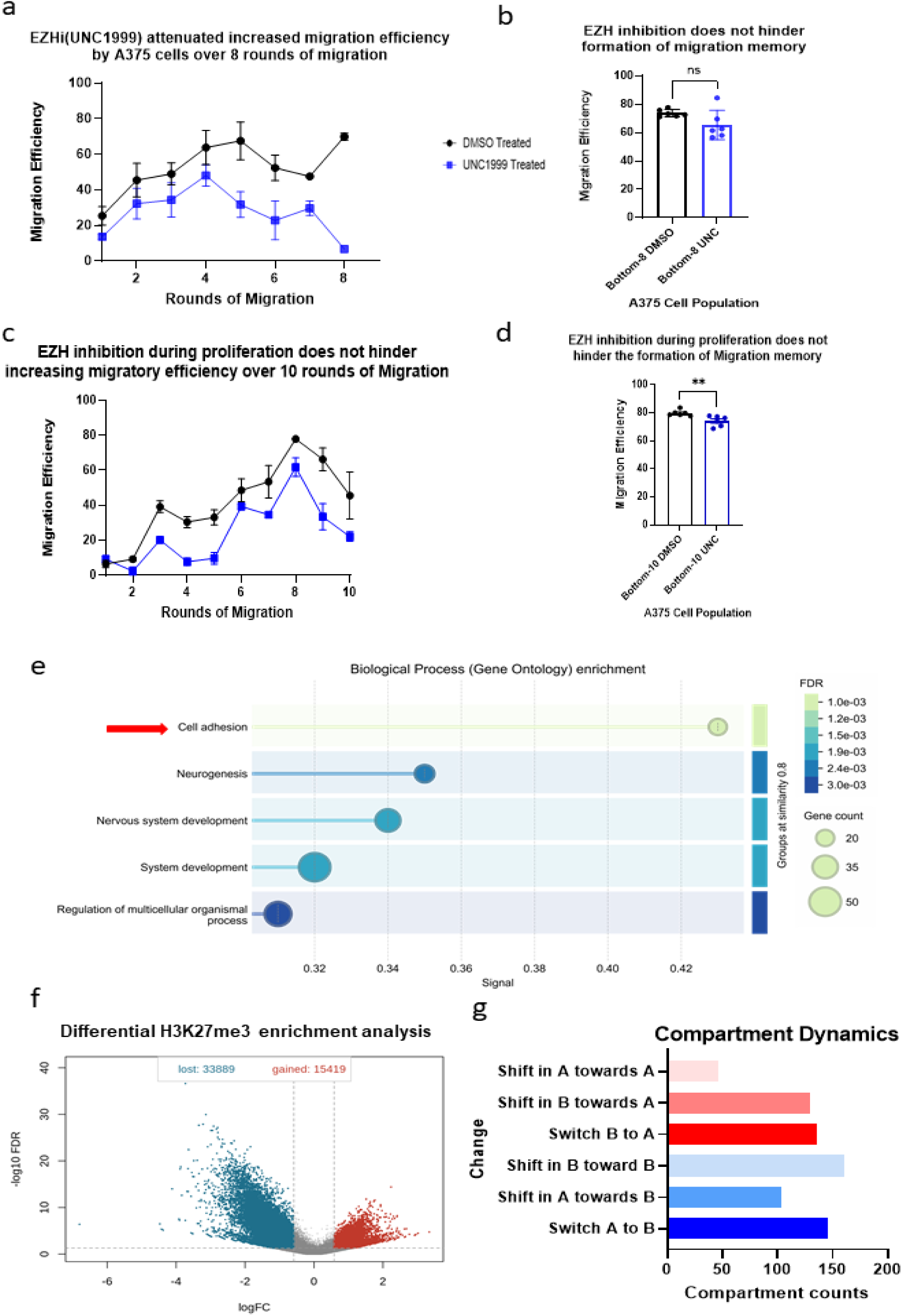
Chronic EZH1 & 2 inhibition led to the selection of a drug-resistant population of A375 melanoma cells with partial similarities to Bottom-10 cells. (A) Line graph showing migration efficiency over successive rounds of constricted migration under UNC1999 treatment during each migration. Each data point represents the mean and standard deviation of two replicates. (B) Bar plot showing the migration efficiency of cells from (A) after 2 passages recovery from treatment. Each bar represents the mean value and standard deviation of three replicates. (C) Line graph showing migration efficiency over successive rounds of constricted migration, with UNC1999 treatment given in between rounds of migration during cell proliferation. Each data point represents the mean value and standard deviation of three replicates. (D) Bar plot showing the migration efficiency of cells after two passages recovery of the cells after the migration shown in (C) Each bar represents the mean value of three replicates. (E). Gene Ontology function enrichment (from StringDB Biological Processes) of genes upregulated in the Bottom-8 A375 cells treated with UNC1999 vs. DMSO during each round of migration. (F) Volcano plot showing differential enrichment of H3K27me3 between chronic UNC1999 treated vs. DMSO treated Bottom-8. (G) Bar plot showing the frequency of genome compartment change types between chronic UNC1999 treated and untreated Bottom-8 cells.

Next, we tested whether the UNC1999 treatment over the rounds of constricted migration inhibited the acquisition of Bottom-10 phenotypes in the Bottom-8 cells that were generated. The generated Bottom-8 cells were challenged in a transwell constricted migration assay after going through two passages without EZH inhibition treatment. Surprisingly, both UNC1999 treated and vehicle treated Bottom-8 cells exhibited comparable migration capabilities, which is similar to Bottom-10 migration (Fig. 7b). This suggests that EZH inhibition does not fully prevent the acquisition of the highly migratory phenotype.

Evidence in the literature has suggested that the epigenome is disrupted and rewritten during cell division^33^; our sequential migration protocol involve proliferation the cells for 48 hours between migrations. Thus, we decided to check if inhibiting the writer of H3K27me3 while the cells are growing in culture in between migrations (but not during migration) impacts their ability to acquire the increased migratory phenotype. As shown in Fig.7c, in comparison to the vehicle-treated cells, UNC1999 treatment in between rounds of migration reduced the rate at which A375 cells acquired increased migratory efficiency over the ten rounds of migration. However, both cell populations continually got better at migration and reached peak migration efficiency at round 8. Next, we checked if this treatment-migration paradigm impacted the migration memory in these cells. UNC1999-treated cells migrated only slightly less efficiently in comparison to the vehicle-treated cells (Fig. 7d).

Next, we assessed transcriptional changes associated with the reduction of migration in the cells over the course of the sequential constricted migration with chronic EZH1&2 inhibition treatment. We found 143 upregulated genes and 97 downregulated genes. Gene ontology analysis of the upregulated genes revealed that, as with acute treatment, cell adhesion is the most significant biological process affected by the set of genes (Fig. 7e). Downregulated genes in these treated cells are involved in biological processes such as Skin development, epithelium development, tissue development (Supp. Fig. 5).

Differential analysis of H3K27me3 enrichment between treated and untreated cells revealed 33,889 loci that had reduced H3K27me3 signal, in contrast to 15,419 loci that gained H3K27me3 (Fig. 7f). This is substantially fewer changed loci, and more balanced between loss and gain, than observed in Bottom-10 cells treated with UNC1999 and migrated only once (Fig. 6d). This suggests that the cells that make it through sequential migration under chronic treatment resist some of the changes normally seen during acute treatment.

To describe the 3D genome changes due to the chronic treatment, we compared the compartment identities and strengths. While the proportion of loci switching compartments is comparable in both directions, loci in the B compartment drifting deeper into the B compartment outnumbers loci in A going deeper towards A (Fig. 7g). Notably, this is the opposite trend than what we observed in acute UNC1999 treatment, where more bins shifted toward A in Bottom-10 cells (Fig 5a).

Overall, continuous treatment with EZH1&2 inhibitors during migration impacts genome structural conformation, epigenomic, and transcriptional regulation in a way that reduces the migratory capability of A375 melanoma. Further, cells that manage to migrate in spite of the treatment to some degree resist these changes.

### Chronic and acute inhibition of EZH1 & 2 resulted in distinct epigenetic effects in A375 melanoma cells

We have presented evidence demonstrating that both acute and chronic inhibition of EZH1&2 led to migratory, transcriptional, epigenomic, and 3D genome structural changes in A375 cells. We finally compared 3D genome compartment dynamics across treatment conditions (Fig 8a,b). We considered all the regions that consistently switched from B to A (Fig. 8c) or A to B (Fig. 8d) with constricted migration for both the originally published Bottom-10 and newly generated vehicle treated Bottom-8 condition. We noted that these genomic regions nearly all experienced a shift in the opposite direction during acute UNC1999 treatment for both Parental and Bottom-10 cell populations. That is, a region that switches from A to B with constricted migration (shown as blue in the first two columns of the heatmap, Fig. 8d) was likely to experience a shift in the A direction during acute UNC1999 treatment (red in the last two columns of the heatmap, Fig. 8d). In contrast, chronic UNC1999 treated Bottom-8 cells (middle column of heatmap) showed an intermediate compartment effect: some bins escaped the effects of acute treatment and match the untreated Bottom-10 compartment shift, while others other regions show a similar effect as acute EZH inhibitor treatment. This analysis suggests that in order to undergo constricted migration, a subset of cells is able to overcome the effects of EZH inhibitor treatment and shift some compartments in the direction typically seen with constricted migration.

**Fig 8:**
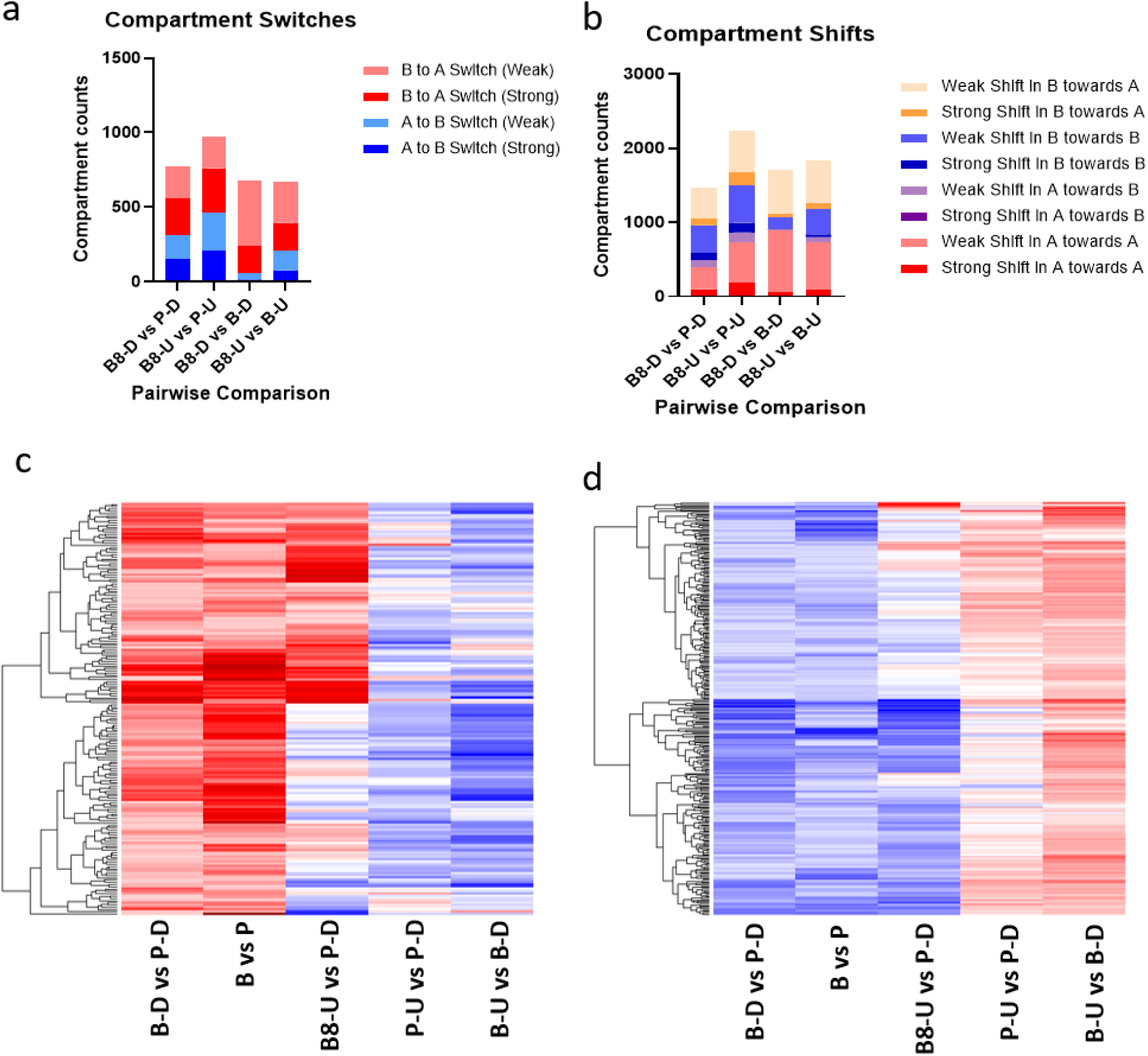
Cells that migrate under chronic EZH inhibition escape some compartment switching effects seen with acute treatment. (A) Stacked bar plot showing the pairwise comparison of compartment switch categories between acute and chronic with migration UNC1999 treatment on A375 cell populations (B) Stacked bar plot showing the pairwise comparison of compartment shifting categories between acute and chronic with migration UNC1999 treatment on A375 cell populations (C and D) Heatmap showing the compartment dynamics in response to acute and chronic with migration UNC1999 treatment of loci previously shown to shift from (C) B to A and (D) A to B between Parental and Bottom-10 cells. (P: Parentals, P-D: Parental DMSO-treated, P-U: Parental UNC1999-treated, B: Bottom-10, B-D: Bottom-10 DMSO-treated, B-U: Bottom-10 UNC1999-treated, B8-U: Bottom-10 Migrated + UNC1999 Treatment)

Overall, evidence presented supports the conclusion that acute and chronic inhibition of EZH1&2 elicits similar, yet distinct effects on melanoma cells. While inhibition of EZH1&2 by UNC1999 does not completely abrogate the acquisition of migratory efficiency in A375 melanoma cells, it prevents the acquisition of the full epigenomic and genome structural conformation seen in Bottom-10 A375 cells, indicating the importance of H3K27me3 in the acquisition and maintenance of constricted-migration-induced epigenetic and 3D genome changes in A375 melanoma cells.

## Discussion

The results presented here support our hypothesis that changes in the epigenetic landscape during successive rounds of migration underlie the 3D genome changes and transcriptional changes we observe in Bottom-10 cells. Earlier evidence in the literature demonstrates that histone enrichment patterns contribute strongly to genome accessibility and overall 3D genome architecture ^29,30^. We extend this finding with the result that histone epigenetic marks can also predict the likelihood of changes in chromatin organization arising from stress. Our results further suggest that H3K27me3 enrichment, which has previously been shown to increase transiently after constricted migration^24^, plays an important role in determining the 3D genome changes in A375 cells after sequential rounds of constricted migration, through its dynamic relationship with loci that switch between the A and B compartments. This argument is further strengthened by relating H3K27me3 and H3K9me3 enrichment to the changes in the transcriptional landscape of Bottom-10 A375 cells that we had previously established. At the genomic locations of genes upregulated in Bottom-10 cells relative to naive parental cells, H3K27me3 enrichment decreased, whereas at genes downregulated in Bottom-10 cells, it increased. At these same locations, however, there were few changes in H3K9me3, suggesting that H3K27me3 plays a larger role in this scenario.

Indeed, overall, our results reveal a strong correlation between H3K27me3 changes and compartment and gene expression changes with migration. Across the body of differentially regulated genes and the loci that switched compartments in the Bottom-10/Parental comparison, H3K27me3 enrichment showed a correlative relationship consistent with the expected behavior of a heterochromatin marker with respect to genome accessibility and transcription. This is consistent with the Polycomb-mediated repression model that has been associated with aggressive metastatic disease^34^. Increased H3K27me3 can silence tumor- and metastasis-suppressive genes, promote cellular plasticity, and support metastatic behavior ^35^, all characteristics associated with A375 Bottom-10 cells. The results presented here also agree with previous evidence demonstrating that increased H3K4me1 enrichment may reflect enhancer priming or reprogramming associated with the acquisition of metastatic competence ^36^. H3K4me1 marks poised and active distal enhancers and, accordingly, metastatic progression in several cancer models has been linked to extensive enhancer remodeling, including the establishment or activation of regions enriched for H3K4me1.

Here we showed that hindering H3K27me3 deposition by inhibiting the catalytic activity of EZH1 and EZH2 reduced the migratory capabilities of both populations of A375 cells, with a more pronounced effect on constricted migration. This raises the possibility that such a treatment modality may be promising both for preventing the initiation of metastasis from a primary tumor and for reducing the likelihood of metastasis from secondary tumors. The evidence presented here further suggests the promise of EZH1 and EZH2 inhibition for oncotherapeutics. Valemetostat, another EZH1 and EZH2 inhibitor, is currently in different phases of clinical trials for several types of malignancies ^37^. Our findings further suggest that, in melanoma metastasis, the therapeutic window for drugs targeting EZH1 and EZH2 lies between the appearance of the tumor and the onset of metastasis. Here, we demonstrated that while acute EZH1 and EZH2 inhibition offers some therapeutic advantage in melanoma generally, naive unmigrated melanoma cells are held stably in their poorly migratory state after just one round of treatment, whereas already proficient migratory cells regain their invasiveness in the absence of continued treatment.

Our results indicate that acute UNC1999 treatment had the most notable effect of decreasing H3K27me3 enrichment at genes and loci that usually switch from A to B and are downregulated during constricted migration. This suggests that the treatment may counteract and prevent some of the increased H3K27me3 at these loci that promote migratory efficiency. Concordantly, acute UNC1999 treatment induced the upregulation of genes involved in cell-to-cell adhesion. This is consistent with evidence in the literature demonstrating that inhibition of EZH2 can alleviate PRC2-dependent repression of epithelial adhesion genes, including CDH1, which encodes E-cadherin ^38,39^. In prostate cancer, H3K27me3 is associated with transcriptional silencing of E-cadherin, resulting in the loss of cell-to-cell adhesion and increased invasive potential ^40^. Consistent with this mechanism, inhibition of the catalytic subunits of PRC2 has been reported to increase E-cadherin expression and reduce metastatic behavior in breast cancer models (Song et al., 2016). These findings suggest that restoration of an epithelial, E-cadherin-mediated cell-to-cell adhesion program may contribute to the anti-invasive effects of EZH1 and EZH2 inhibition in melanoma and other tumor types.

Consistent with the effect of the EZH1/2 inhibitor valemetostat in a small-cell lung cancer model,^37^ UNC1999 induced reorganization of the 3D genome structure in our system. Bottom-10 cells, a distinct population, underwent far more compartment switches toward the active A compartment, whereas parental cells switched primarily in the opposite direction. This result further suggests that, although both cell populations showed a similar migration-related response to treatment, the underlying pathways are largely distinct and may be associated with the cells’ ab initio higher-order genome conformation. The partial reversal of chromosomal compartment identity at loci in Bottom-10 cells after UNC1999 treatment implies that genome structure, beyond gene function, plays an important role in maintaining migration efficiency. Previous evidence has suggested that the heterochromatin-to-euchromatin ratio of the nucleus correlates with nuclear stiffness and pliability ^41^, which in turn affects the ability of cells to pass through tight spaces. Our results suggest that reducing nuclear H3K27me3 content may increase nuclear flexibility beyond the critical point that provides the intranuclear pressure required for efficient migration during metastasis. Further work is needed to confirm this proposed relationship between treatment and nuclear mechanics in our model system.

The heterogeneity of cancer cell populations implies the presence of cell subtypes, including those resistant to various therapeutic compounds. Our results indicate that, although UNC1999 treatment altered H3K27me3 enrichment across the genome, the cells that were still able to migrate in the presence of treatment were more likely to resist these treatment-induced epigenetic changes. We also demonstrated the distinct and combined effects of migration and UNC1999 treatment on H3K27me3 enrichment in our model system, establishing a non-stochastic relationship between chemical and mechanical influences on genomic H3K27me3 enrichment in our model.

The results presented here suggest that UNC1999 treatment resistant cells are less affected by both acute and chronic treatment and can repopulate a highly migratory population of A375 cells. Interestingly, treating cells before and during migration produced a more pronounced reduction in migration efficiency than treating between migrations, when the cells are thought to be breaking down and reassembling their epigenetic landscape ^42–44^. Earlier evidence has proposed either the acquisition of mutations or an EMT-associated drug-tolerant persister state as the mechanism by which cancer cells evade treatment ^45,46^. Our previous work suggested that Bottom-10 A375 cells acquire transcriptional and morphological characteristics indicative of epithelial-mesenchymal transition ^6^. This again reiterates the need for combination or adjuvant strategies in oncotherapeutics.

Our results demonstrate that while acute UNC1999 treatment resulted in a genome-wide loss of H3K27me3 enrichment, chronic treatment during sequential migration resulted in a less dramatic loss and a genome compartment profile that leaned largely toward the inactive heterochromatin compartment. The chronic model likely represents a simultaneous pharmacologically induced reduction and mechanically induced increase in H3K27me3 enrichment. The gain of H3K27me3 at some loci and the overall shift of the genome toward a more heterochromatinized state suggest that mechanical induction of heterochromatinization outweighs pharmacological removal and is selected for in cells that are able to migrate even in the presence of the drug. Importantly, the findings presented here demonstrate that although acute EZH1/2 inhibition was able to shift chromatin compartment switches observed in Bottom-10 cells back toward their initial identities in parental cells, chronic inhibition during sequential migration prevented only part of the 3D genome changes seen in the Bottom-10 population. This suggests that some compartment changes observed in Bottom-10 cells are dispensable for achieving a final high migratory efficiency, but others are necessary.

In conclusion, the evidence presented here supports the argument that H3K27me3 plays a key role in encoding and maintaining higher migratory efficiency in sequentially constricted A375 melanoma cells, and that it does so through transcriptional regulation of migration-associated genes and reorganization of the overall physical structure of the genome.

## METHODS AND PROTOCOLS

### KEY RESOURCES TABLE

| Num | REAGENT | SOURCE | IDENTIFIER |
| --- | --- | --- | --- |
| 1 | A375 human melanoma cell line | ATCC | CRL-1619 |
| 2 | DMEM | Corning | 10-013-CV |
| 3 | Fetal bovine serum (FBS) | Corning | 35-010-CV |
| 4 | Penicillin–streptomycin | Gibco | 15140-122 |
| 5 | L-glutamine | Gibco | 25030-081 |
| 6 | CUTANA CUT&RUN Kit | EpiCypher | 14-1048-48rxn |
| 7 | CUTANA pAG-MNase | EpiCypher | 15-1016 |
| 8 | DNA Purification beads | EpiCypher | 21-1407k |
| 9 | H3K27me3 antibody | EpiCypher | 13-0055 |
| 10 | H3K4me1 antibody | EpiCypher | 13-0057 |
| 11 | H3K27ac antibody | EpiCypher | 13-0059 |
| 12 | H3K9me3 antibody | Abcam | ab176916 |
| 13 | H3K27me3 antibody (IF) | Abcam | Ab6002 |
| 14 | Monarch PCR & DNA Cleanup Kit | New England Biolabs | 13-0041 |
| 15 | NEBNext Ultra II DNA Library Prep Kit | New England Biolabs | (Arima protocol A16041v01) |
| 16 | Alexa Fluor 488 secondary antibody | Thermo Fisher | R37117 |
| 17 | Alexa Fluor 594 secondary antibody | Thermo Fisher | R37120 |
| 18 | DAPI | Thermo Fisher | 62248 |
| 19 | Arima-HiC+ Kit | Arima Genomics | A160134 v01 |
| 20 | 5 µm Transwell filters | VWR | 10769-236 |
| 21 | 12 µm Transwell filters | VWR | 10769-224 |
| 22 | 5 µm pore membrane (custom 10 cm inserts) | Sterlitech | PCT5014220 |

### SOFTWARE AND ALGORITHMS

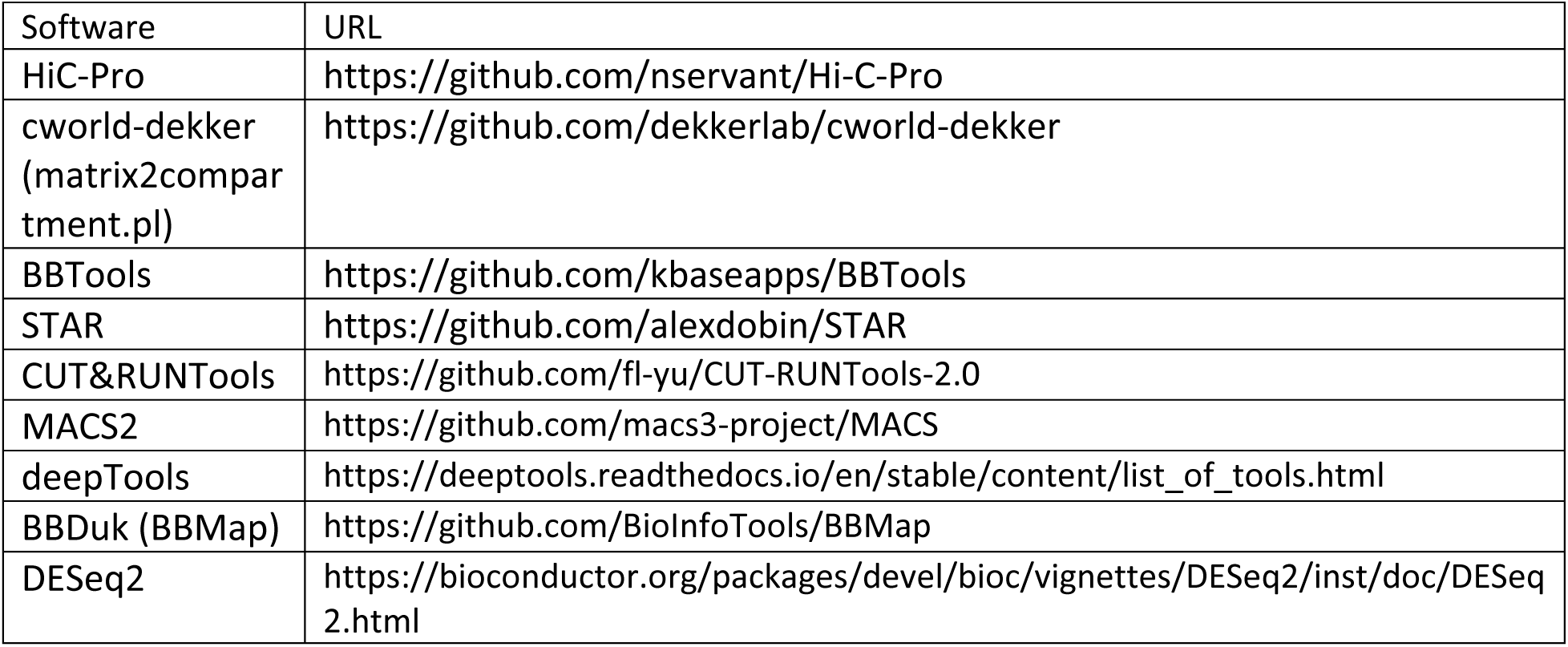
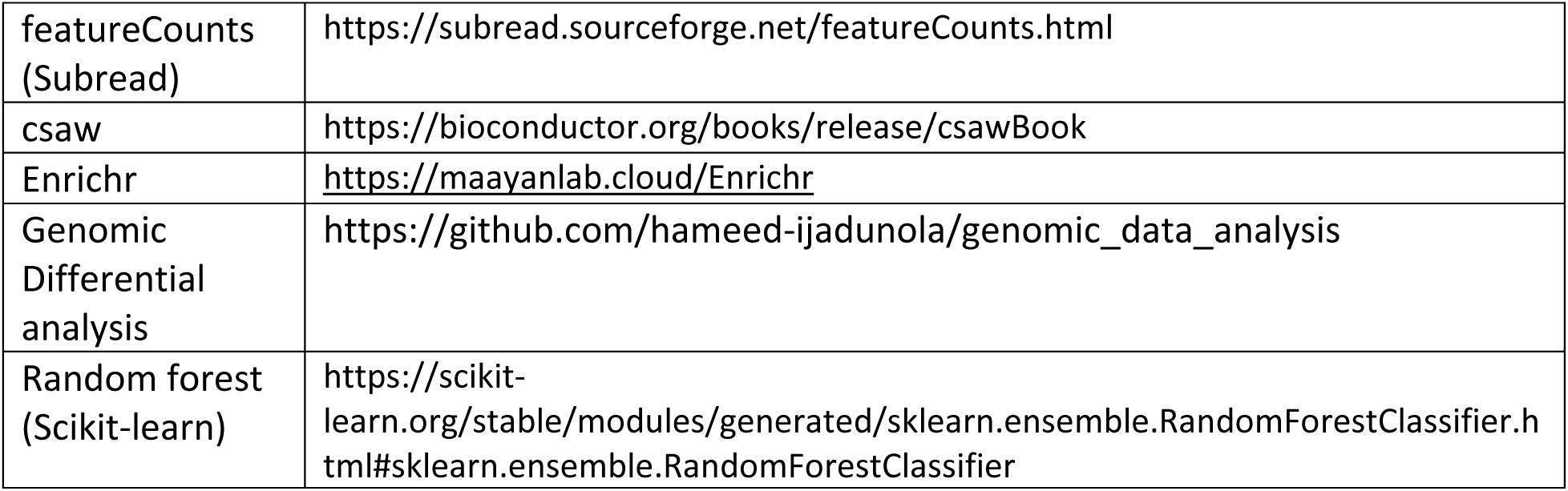

### A375 CELL CULTURE

A375 human melanoma cells (CRL-1619) were obtained from ATCC. Cells were verified to be mycoplasma-negative before use. Cells were cultured in complete DMEM (Corning 10-013-CV) supplemented with 10% fetal bovine serum (FBS), 1% penicillin–streptomycin, and 1% L-glutamine at 37°C in a humidified incubator with 5% CO2. Parental cells were defined as those taken directly from this initial culture. Bottom-10 cells were previously migrated through 10 rounds of 5-micron Transwell pores.

### IMMUNOFLUORESCENCE STAINING AND CONFOCAL IMAGING

Approximately 25,000 cells from each A375 subpopulation were seeded into 35-mm MatTek dishes containing poly-D-lysine–treated coverslips and allowed to attach overnight. Media was replaced with fully supplemented media containing either an EZH inhibitor or DMSO vehicle and incubated for 24 h, followed by incubation in unsupplemented media containing the same treatment for an additional 24 h (to mimic Transwell migration conditions). Cells were fixed with 4% formaldehyde for 10 min and washed three times with PBS (5 min each). Cells were permeabilized in permeabilization buffer (10% goat serum, 0.5% Triton X-100 in PBS) for 1 h at room temperature.

Primary antibodies (H3K27me3, Abcam ab6002; H3K9me3, Abcam ab176916) diluted in antibody dilution buffer (5% goat serum, 0.25% Triton X-100 in PBS) were applied and incubated overnight at 4°C. Cells were washed three times with PBS (5 min each) and incubated with secondary antibodies (Alexa Fluor 488 and Alexa Fluor 594; Thermo Fisher R37117 and R37120; 2 drops per 1 mL PBS) for 30 min at room temperature. Cells were washed three times with PBS and stained with DAPI (Thermo Fisher 62248) for 15 min at room temperature. Coverslips were mounted on slides using Fluoromount-G, sealed, and cured in mounting medium for 24 h before imaging. Imaging was performed on a Leica SP8 confocal microscope using a 63× oil-immersion objective.

### TRANSWELL MIGRATION ASSAYS

Transwell filters with 5-µm pores (VWR 10769-236) and 12-µm pores (VWR 10769-224) were used. The underside of each filter was coated with 40 µL fibronectin (10 µg/mL) for ∼40 min. For each well of a 24-well plate, 500 µL 1× DMEM (Corning) with full supplements was added to the bottom chamber. A375 cells were detached at 80–90% confluency and resuspended in unsupplemented 1× DMEM to 1 × 10^5 cells per 100 µL. Transwells were placed into wells, and 100 µL of cell suspension was added to the top chamber. Cells were incubated for 24 h at 37°C and 5% CO2.

Migration efficiency was quantified as previously described by Golloshi et al^6^. Cells from the top and bottom chambers were collected separately; remaining adherent cells were detached with trypsin, and recovered cells were counted to calculate percent migration as #bottom/(#top + #bottom). For sequential constricted migration, bottom-chamber cells were expanded for 48 h prior to the next migration round.

### TRANSWELL MIGRATION ASSAYS WITH EZH INHIBITORS

For inhibitor experiments, cells were pre-treated with 5 μM of EZH inhibitors (or 0.005% DMSO) in fully supplemented 1× DMEM (Corning) for 24 h prior to plating into Transwells. Bottom chambers were prepared using fully supplemented 1× DMEM containing the EZH inhibitor. Cells were detached from treated cultures and seeded into the upper chamber at 1 × 10^5 cells per 100 µL in unsupplemented 1× DMEM containing the EZH inhibitor. For matched conditions where cells were treated with EZH inhibitors but not migrated, cells were treated with inhibitor for 24 h in fully supplemented media and then for 24 h in unsupplemented media with inhibitors to match the media conditions of the migration experiment.

### TRANSWELL MIGRATION ASSAYS USING 10-CM DISH INSERTS

Custom 10-cm Transwell inserts adapted from Playter et al. ^47^were used. A 5-µm pore membrane (Sterlitech PCT5014220) was attached to the bottom of each insert using a two-part epoxy applied with a P1000 pipette tip. Epoxy was cured for 24 h at room temperature, and excess membrane was trimmed. Inserts were sprayed with 70% ethanol and sterilized under UV in a biosafety cabinet for ∼2 h. Filters were coated with fibronectin by placing 1 mL fibronectin (10 µg/mL) on parafilm and placing filters on top for ∼40 min; excess was removed prior to migration assays. Migration assays were performed as above (with or without EZH inhibitors) with adjusted inputs: 15 million cells were added, with 7 mL and 9 mL in the top and bottom chambers, respectively. For incubation, the apparatus was placed inside a 15-cm dish to maintain sterility. For sequential constricted migration, bottom-chamber cells were expanded in a T-75 flask for 48 h prior to subsequent rounds.

### CLEAVAGE UNDER TARGET AND RELEASE USING NUCLEASE (CUT&RUN) AND LIBRARY PREPARATION

CUT&RUN was performed for histone-mark profiling using the EpiCypher CUTANA CUT&RUN protocol (Version, 2022) with minor modifications. Approximately 5 × 10^5 A375 cells were washed twice in Wash Buffer and resuspended in 100 µL Wash Buffer. Activated concanavalin A beads (10 µL) were added, and samples were incubated for 10 min at room temperature to bind cells to beads. Beads were collected on a magnetic rack, and the supernatant was discarded. Beads were resuspended in 50 µL cold Antibody Buffer to form a slurry. The K-MetStat panel (2 µL) was added and mixed, after which 1 µL ChIP-grade antibody was added and mixed by gentle pipetting. Antibodies used were: H3K27me3 (EpiCypher 13-0055), H3K4me1 (EpiCypher 13-0057), H3K27ac (EpiCypher 13-0059), and H3K9me3 (Abcam ab176916). Samples were incubated overnight at 4°C on a nutator.

On day 2, beads were collected on a magnetic rack, the supernatant was removed, and beads were washed twice with 200 µL ice-cold Cell Permeabilization Buffer containing 0.05% digitonin. Beads were resuspended in 50 µL of the same buffer, and 2.5 µL CUTANA pAG-MNase was added to each sample and mixed by gentle pipetting. Samples were incubated for 10 min at room temperature. Beads were then collected and washed twice with 200 µL ice-cold Cell Permeabilization Buffer containing 0.05% digitonin, and resuspended in 50 µL of the same buffer. To activate MNase digestion, 1 µL of 100 mM CaCl2 was added to each sample, mixed gently, and incubated for 2 h at 4°C on a nutator. Reactions were stopped by adding 33 µL stop master mix (Stop Buffer plus 1 µL spike-in E. coli DNA) and incubating samples for 10 min at 37°C in a preheated thermocycler. Beads were placed on a magnetic rack, and released CUT&RUN DNA was transferred to new 1.5 mL tubes. DNA was purified using the Monarch PCR & DNA Cleanup Kit (New England Biolabs 13-0041) according to the manufacturer’s instructions. Purified DNA was stored at −20°C overnight.

On day 3, 5 ng purified CUT&RUN DNA (in 25 µL 0.1× TE) was used to generate sequencing libraries using the CUT&RUN Library Prep Reactions Kit according to the manufacturer’s instructions. DNA ends were repaired by adding 5 µL end-repair master mix (4.2 µL End Prep Buffer + 1.8 µL End Prep Enzyme) and running the recommended thermocycler program. Samples were then placed on ice in pre-chilled aluminum blocks. Adapter ligation was performed by adding 1.25 µL of 1.5 µM Illumina adapter and 15.5 µL ligation master mix (16.5 µL ligation mix + 0.55 µL ligation enhancer), mixing gently, and incubating for 15 min at 20°C. Next, 1 µL U-excision enzyme was added, and samples were incubated for 15 min at 37°C to remove uracil-containing hairpins. Libraries were purified using SPRIselect beads according to the manufacturer’s instructions. Libraries were amplified using unique index pairs with 13 PCR cycles. Final library quality and quantity were assessed using an Agilent Bioanalyzer. Libraries were sequenced by GeneWiz using 150 bp paired-end reads to a minimum depth of 10 million reads per sample.

### CUT&RUN DATA PROCESSING

Paired-end 150 bp reads were processed with CUT&RUNTools v2.1^48^. TruSeq3 adapters were trimmed with Trimmomatic v0.36^49^ (`ILLUMINACLIP:2:15:4:4:true LEADING:20 TRAILING:20 SLIDINGWINDOW:4:15 MINLEN:25`), followed by the kseq step that removes residual adapters from the short fragments. Trimmed reads were aligned to hg19 with Bowtie2^50^ using `--very-sensitive-local --phred33 -I 10 -X 700`. Alignments were sorted and duplicates removed with Picard v2.8.0^51^. Peaks were called on deduplicated alignments with MACS2^52^ in paired-end mode (`-f BAMPE -g hs --broad --broad- cutoff 0.1 --keep-dup all`). Coverage tracks were generated with deepTools `bamCoverage`^31^ at 10-bp resolution with CPM normalization. Reads were additionally aligned to the *Escherichia coli* K-12 DH10B genome to quantify spike in *E. coli* DNA.

### CUT&RUN GENOMIC DIFFERENTIAL ANALYSIS

Differential histone modification signal between the control and test groups was computed on a standalone Jupyter notebook (https://github.com/hameed-ijadunola/genomic_data_analysis) built using pyBigWig (https://github.com/deeptools/pyBigWig). We binned each CUT&RUN CPM normalized bigWig file at 5 kb by taking the mean signal within each bin; bins with no coverage were treated as zero. For each pair of conditions (e.g., Parental and Bottom-10), subtractions were performed at each bin in the genome for each possible pair of replicates (P-R1 vs. B10-R2, P-R1 vs. B10-R1, P-R2 vs. B10-R1, P-R2 vs. B10-R2) and then the differences were averaged at each bin to yield a pairwise mean difference track. A per-bin threshold was computed using these replicate comparisons as 2 standard deviations of the mean pairwise differences in that bin and bins whose mean pairwise difference did not exceed the threshold in either direction was excluded from further comparison. The differences per bin were then used to identify the top 1000 changing regions for visualization and clustering in Figure 6e.

### CUT&RUN STATISTICAL ANALYSIS CUT&RUN

differential binding was assessed with csaw v1.40.0^53^, which counts reads in sliding windows and requires no prior peak calling. Reads were counted in 2-kb windows spaced 500 bp apart, with duplicates removed and a maximum fragment length of 700 bp. Windows were retained only if enrichment over the genomic background, which was estimated from 10-kb bins genome-wide, exceeded 3-fold (log₂ = 1.585). Where multiple window sizes were used, results were merged into shared regions with `mergeWindowsList` and combined with `combineTests`, so each region carries a single region-level FDR. Counts were modeled in edgeR v4.4.0^54^ using a negative binomial generalized linear model; dispersions were estimated with `estimateDisp` and significance was tested by quasi-likelihood F-test (`glmQLFit` with robust estimation, followed by `glmQLFTest`). Regions with FDR < 0.05 were considered differentially bound.

Since EZH inhibitor treatments can cause major changes in the global levels of H3K27me3, we tested methods for adjusting CUT&RUN signals to account for global enrichment differences. We first tried to use reads mapped to the *E. coli* spike-in DNA, but the read counts varied according to both technical experimental variations and biological conditions, and counts were often not high enough to allow robust quantification, as has been previously reported in the literature^55^. This could be due to inconsistent amounts of spike-in material in all samples. As a result, spike-in scaling was not applied. Instead, we normalized against genomic background, applying TMM to 10-kb bins (`csaw::normFactors`).

### CROSSLINKING CELLS FOR HI-C

Approximately 5–10 million A375 cells were crosslinked in 5 mL HBSS containing 1% formaldehyde for 10 min at room temperature with gentle shaking. Crosslinking was quenched by adding 277 µL of 2.5 M glycine and mixing. Cells were pelleted in a cooled centrifuge and supernatant was removed. Pellets were washed by resuspending in 5 mL ice-cold HBSS containing 50 µL protease inhibitor, followed by centrifugation and complete removal of supernatant. The crosslinked pellet was flash-frozen in liquid nitrogen and stored at −80°C.

### HI-C EXPERIMENTS, SEQUENCING AND DATA PROCESSING

Hi-C was performed using the Arima-HiC+ Kit (Arima Genomics) following the Mammalian Cell Lines protocol (A160134 v01). Libraries were prepared following Arima recommendations for the NEBNext Ultra II DNA Library Prep Kit (protocol version A16041v01). Sequencing was performed by GeneWiz on an Illumina NovaSeq using 150 bp paired-end reads. Reads were mapped to hg19, filtered, and iteratively corrected using HiC-Pro.

Compartment analysis was performed using principal component analysis with the matrix2compartment.pl script in the cworld-dekker pipeline. Compartment identity was assigned using PC1 values from 250-kb binned matrices: using the correlation with gene density (highest gene density regions are nearly always in the A compartment and lowest gene density regions are nearly always in the B compartment), the sign of the PC1 profile was assigned so that PC1 > 0 indicated compartment A and PC1 < 0 indicated compartment B. Compartment switches between two groups were defined as sign changes in PC1 (A to B or B to A). Compartment shifts were defined as PC1 differences > 0.02 (toward A) or < −0.02 (toward B), corresponding to a 10% change in compartment score. Strong shifts were defined as PC1 differences > 0.04 or < −0.04, corresponding to a 20% compartment score change.

### RNA-SEQ EXPERIMENTS

RNA was extracted using the Qiagen RNeasy Plus Mini Kit. Approximately 3 × 10^5 to 7 × 10^5 cells were lysed and homogenized, and genomic DNA was removed using gDNA Eliminator columns. Samples were washed with ethanol, and total RNA was eluted. RNA quantity and quality were assessed by NanoDrop. RNA-seq libraries were prepared using a poly(A) mRNA approach and sequenced by GeneWiz.

### RNA-SEQ ANALYSIS

Adapters were trimmed with BBDuk (BBMap v38.18^56^; `ktrim=r k=23 mink=11 hdist=1`), followed by 3′ quality trimming at Q28 (`QTRIM=R TRIMQ=28`). Trimmed reads were aligned to GRCh38, GENCODE v49 with STAR v2.7.6a^57^ using `--outFilterScoreMinOverLread 0.2 --outFilterMatchNminOverLread 0.2`. Alignments were coordinate-sorted and indexed with SAMtools v1.16.1^58^, and gene-level counts generated with featureCounts^59^ (Subread v2.1.1) over exons grouped by `gene_id`, counting fragments (`-p -- countReadPairs`). CPM-normalized coverage tracks were generated with deepTools^31^ (v 3.5.5) bamCoverage at 10-bp resolution for visualization only. Differential expression was assessed with DESeq2 v1.46.0^60^ in R v4.4.3. Genes were retained if they had at least 10 counts in at least 2 replicates of either condition. P values were adjusted by the Benjamini-Hochberg procedure with independent filtering, and genes with adjusted P < 0.05 were considered differentially expressed. Shrunken fold changes (apeglm) were used for ranking and visualization only. To link gene expression data with CUT&RUN and Hi-C data, we linked the log2fold change for each gene with the annotated coordinates of that gene in hg19. A gene was considered overlapping with a 250 kb bin compartment switch if the transcription start site fell within the bin.

### GENE FUNCTION ENRICHMENT ANALYSES

Differentially expressed genes (DEGs) identified by DESeq2 were analyzed for functional enrichment using STRING-db^61^ and Enrichr^62^. STRING-db was used to identify enriched Gene Ontology (GO) Biological Process terms, and Enrichr was used to assess GO Molecular Function enrichment. Enrichment significance was evaluated using multiple-testing–adjusted p-values (false discovery rate, FDR). Terms meeting the predefined threshold (e.g., FDR < 0.05) were considered significant.

### PREDICTING COMPARTMENT SWITCHING FROM PARENTAL CHROMATIN STATE

To predict compartment switching from histone marks, we extracted the mean CPM-normalized CUT&RUN signal of the four marks (H3K27me3, H3K27ac, H3K9me3, and H3K4me1) from parental cell bigWig files in 250 kb bins (Data S1). Based on our previously published Hi-C data, we classified 250 kb bins as switching, shifting, or stable (see Data S1) for compartment change classifications). Bins are the unit of analysis, so each label corresponds to a single observation. For classification, we used random forest models in scikit-learn^63^ (v1.8.0) with 500 trees, a maximum depth of 12, and class_weight=“balanced_subsample” to offset the imbalance between switching and stable bins. All other parameters were left at their defaults. Model performance was assessed by five-fold stratified cross-validation and reported as ROC-AUC and precision-recall AUC. XGBoost was run alongside as a check and gave comparable results, so random forest is reported throughout. Because neighboring bins tend to share compartment and epigenomic state, random folds can place a bin and its neighbor on opposite sides of a split. We therefore repeated the cross-validation holding out whole chromosomes (`StratifiedGroupKFold` grouped by chromosome).

To confirm that the model learned from epigenomic state rather than starting compartment, we compared models trained on the marks alone, parental PC1 alone, both together, and marks shuffled across bins. To assess significance, class labels were shuffled 2,000 times and the model refitted on each shuffle; the observed AUC was compared against this empirical null and a permutation P value computed using the (+1) correction of Phipson and Smyth (2010)^64^. Feature contributions were quantified using SHAP values (TreeExplainer).

### OMICS INTEGRATION

Per-bin mark changes were computed as log₂ ((Bottom10 + ε) / (Parental + ε)). A separate ε was used for each histone mark and defined as 1% of the median signal across all bins from both conditions for that mark (H3K27me3, 1.63 × 10^−4; H3K27ac, 1.91 × 10^−4; H3K4me1, 2.04 × 10^−4; H3K9me3, 2.31 × 10^−4). This choice keeps low-signal bins finite while remaining below the minimum observed signal (9.2 × 10^−4), avoiding compression of ratios. Only migration-axis contrasts (Parental vs Bottom10) were utilized for integration. Bins were deduplicated by coordinates such that each bin contributed one value per mark (n = 10,576 bins with both compartment calls and CUT&RUN signal). Distributions across the four compartment-switch categories were compared using a Kruskal-Wallis test followed by Dunn’s test with Benjamini-Hochberg correction (six comparisons per mark), and Cliff’s delta was used as an effect size. Analyses were performed in Python v3.11.15 using SciPy v1.17.1 and scikit-posthocs v0.14.0. All analyses were conducted at the genomic-bin level; each 250-kb bin appears exactly once per test to avoid pseudo-replication that would occur if genes within bins were treated as independent observations.

For the three-way integration, genes were assigned to 250-kb bins by transcription start site. Since a bin can contain several genes, gene-level log2 fold changes were aggregated to a single value per bin by taking the mean across the genes it contained, and bins with no mapped gene were excluded. This keeps the unit of analysis consistent across all three modalities and avoids inflating the sample size by treating genes that share a bin as independent. The association between mark change and expression change was quantified by Spearman correlation. To distinguish a direct relationship from one arising because both quantities follow the compartment switch, we also computed the partial Spearman correlation conditioning on ΔPC1, obtained by rank-transforming all three variables and correlating the residuals of mark change and expression change after regression on ΔPC1.

### QUANTIFICATION AND STATISTICAL ANALYSIS

Unless otherwise indicated, statistical analyses were performed in GraphPad Prism. Statistical details (sample size, tests used, and definitions of center and dispersion) are provided in the figure legends.

## Supplementary Information

File contains 5 supplementary figures

## Acknowledgements

We thank all members of the McCord lab for helpful discussion and feedback. We acknowledge the UTK Advanced Microscopy and Imaging Center and Genomics Core for assistance with confocal microscopy and sequencing.

## Competing Interests

The authors declare no competing interests.

## Funding

This work was supported by the National Institutes of Health [NIGMS grant R35GM133557 to R.P.M]. C.R. was funded by an NSF Summer Undergraduate Research Experience award.

## Notes

### Competing Interest Statement

The authors have declared no competing interest.

