## Supplemental Figures for "H3K27me3 Drives Constricted Migration–Induced 3D Genome Rewiring in Melanoma"

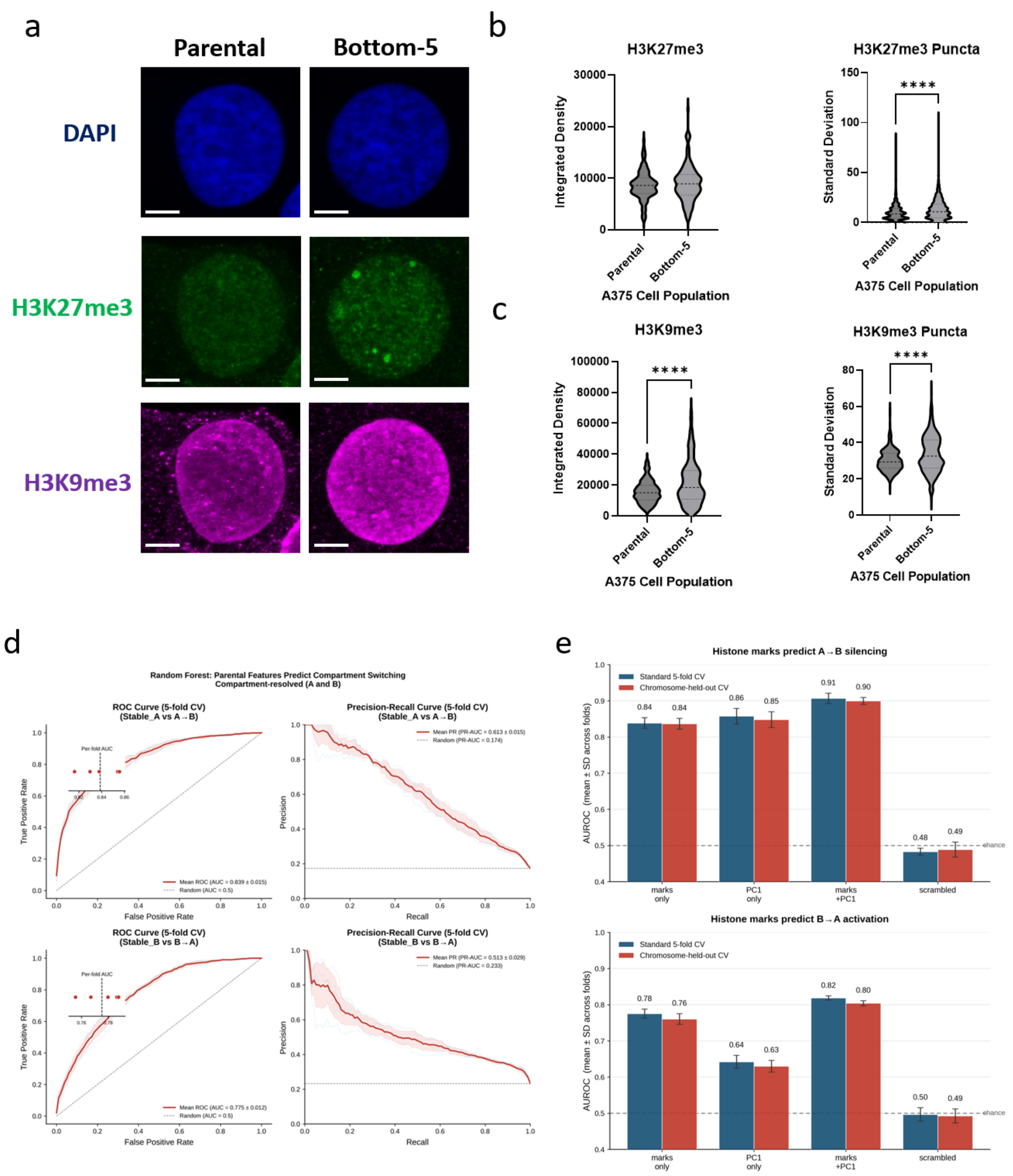


**Supplementary Fig 1: Visible changes in histone marks after constricted migration and additional results from the model predicting compartment switches from histone mark initial state.** (a) Representative images showing Parental and Bottom-5 cells (scalebar = 5 microns). (b) Quantification of integrated densities and standard deviation of H3K27me3 signals in the A375 cell population. Median and quartiles shown with dashed lines. (c) Quantification plots showing the integrated densities and standard deviation of H3K9me3 signals in the A375 cell population. A minimum of 120 nuclei from at least 3 slide images were quantified. (**** p<0.0001, Welch's t test) (d) Plots showing the accuracy of prediction and precision and recall of the compartment changes in the Bottom-10 cells with the Random Forest histone modification model. (e) Bar plots showing the comparison of the accuracy of prediction using enrichment patterns of histone epigenetic markers, Hi-C compartment data, or a combination of both by a Random Forest model.

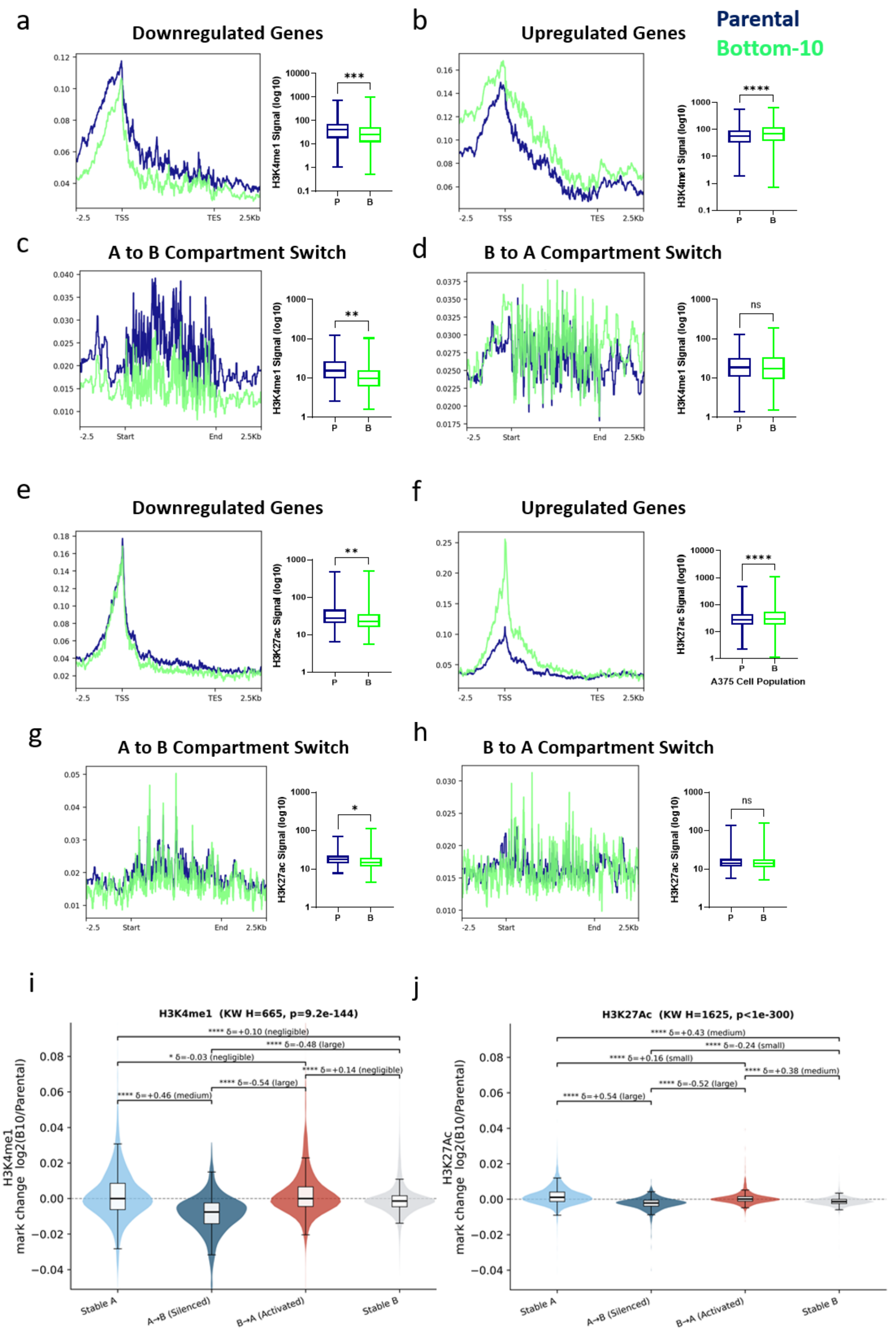


**Supplementary Fig. 2: H3K4me1 and H3K27Ac enrichment levels compared with transcriptional changes and compartment switches in A375 melanoma cells** (a-h) Pile-up plots show the average histone modification signal in Parental (blue) or Bottom-10 (green) cells across sets of differentially regulated genes scaled from transcription start to transcription end (a, b, e, f) or across sets of 250 kb bins that switch compartments (c, d, g, h). H3K4me1 signals are shown in a-d while H3K27ac signals are shown in e-h. Paired boxplots show the distribution of signals across the sets of regions averaged in the pileup. Each point that contributes to the boxplot is the average signal across one gene or one 250 kb bin. Boxplots show minimum to maximum range, quartiles, and median. Significance calculated by Welch's t test (* $p<0.05$, ** $p<0.01$, *** $p<0.001$, **** $p<0.0001$) (i) Violin plot showing log2 fold change in H3K4me1 signal from Parental to Bottom-10 cells across 250 kb bins categorized according to whether they show stable or altered compartment state between Parental and Bottom-10 cells. The significance (stars as described above) is calculated using Kruskal Wallis and the effect size is calculated using Cliff's delta. (j) same as (i) but for H3K27Ac signals along the genome.

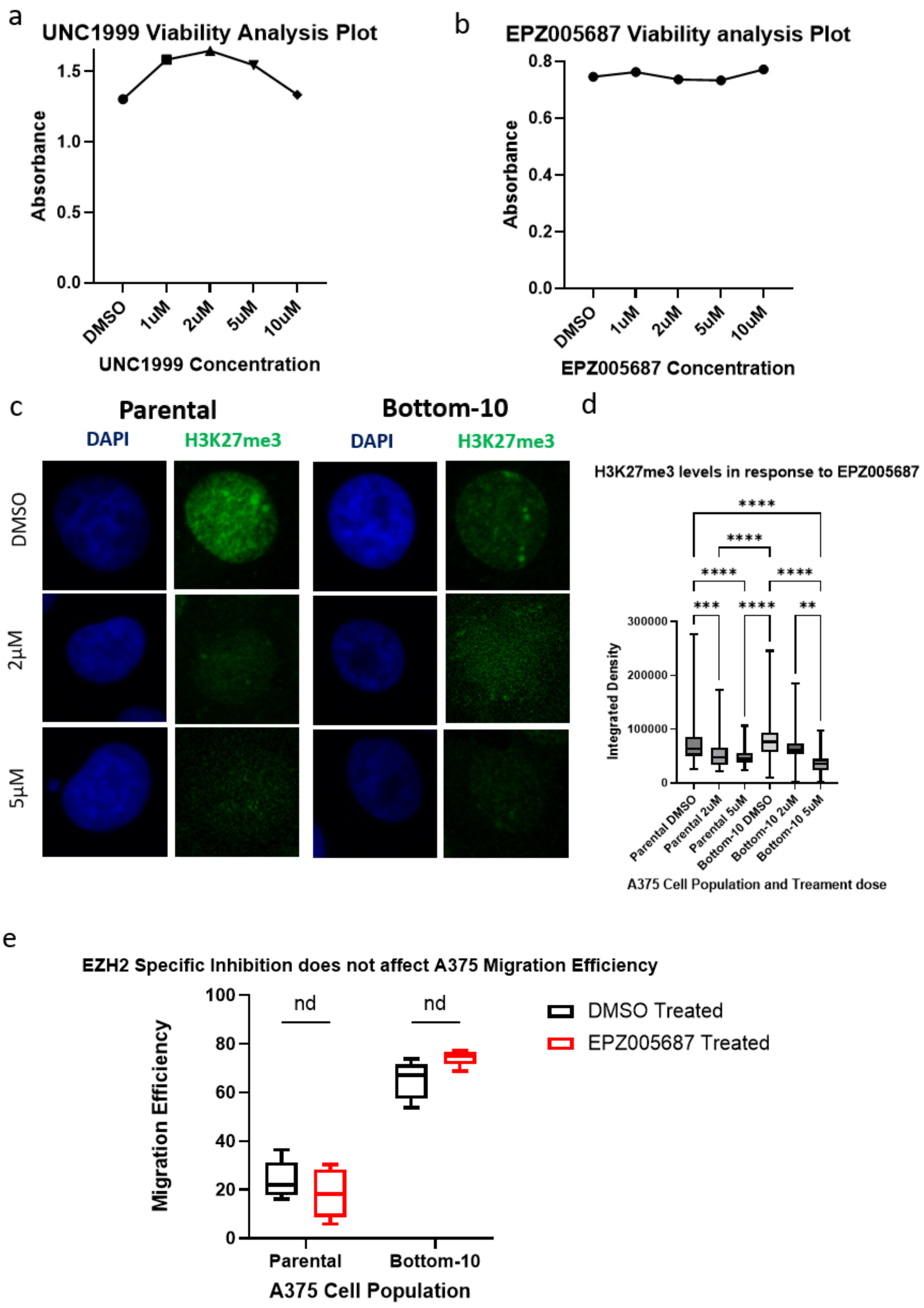


**Supplementary Fig. 3: EZH2-specific inhibition significantly reduces H3K27me3 levels without affecting the migration phenotype of A375 cells.** (a) Line graph showing the mean absorbance values of the MTS assay at different concentrations of UNC1999 treatment in A375 cells. (b) Line graph showing the mean absorbance values of the MTS assay at different concentrations of EPZ005687 treatment in A375 cells. (c)

Representative fluorescent image showing Parental and Bottom-10 cells treated with increasing doses of EPZ005687. (d) Quantification plots show the integrated density of H3K27me3 signals in both the parental and Bottom-10 A375 cell populations after 48 h treatment with EPZ0005687. We quantified at least 120 nuclei from at least 3 slide images. P values refer to an ANOVA test. (*** p<0.001, **** p<0.0001) (e) Box plot showing migration efficiency of A375 cell populations under 5 µM EPZ0005687 treatment. Each box represents 8 replicates. “nd” means t-test p-value > 0.05.

a

| GO Term | P-value | Adjusted P-value | Odds Ratio | Combined Score |
|---|---|---|---|---|
| Laminin-1 Binding (GO:0043237) | 0.002498 | 0.031863 | 555.1667 | 3326.771 |
| Integrin Binding Involved in Cell-Matrix Adhesion (GO:0098640) | 0.002498 | 0.031863 | 555.1667 | 3326.771 |
| Nerve Growth Factor Binding (GO:0048406) | 0.002498 | 0.031863 | 555.1667 | 3326.771 |
| Interleukin-1 Binding (GO:0019966) | 0.003994 | 0.031863 | 317.1905 | 1751.859 |
| Neurotrophin Binding (GO:0043121) | 0.003994 | 0.031863 | 317.1905 | 1751.859 |
| Cell-matrix Adhesion Mediator Activity (GO:0098634) | 0.005488 | 0.031863 | 222 | 1155.57 |
| Lipoprotein Lipase Activity (GO:0004465) | 0.006482 | 0.031863 | 184.9815 | 932.0597 |
| Death Receptor Activity (GO:0005035) | 0.006482 | 0.031863 | 184.9815 | 932.0597 |
| Heparan Sulfate Proteoglycan Binding (GO:0043395) | 0.006979 | 0.031863 | 170.7436 | 847.7047 |

b

| GO Term | P-value | Adjusted P-value | Odds Ratio | Combined Score |
|---|---|---|---|---|
| MHC Class II Protein Complex Binding (GO:0023026) | 3.50E-05 | 0.013581 | 9.495845 | 97.41683 |
| Serine-type Peptidase Activity (GO:0008236) | 5.18E-05 | 0.013581 | 3.177849 | 31.35712 |
| Serine-type Endopeptidase Activity (GO:0004252) | 7.79E-05 | 0.013611 | 3.336912 | 31.56627 |
| Receptor Ligand Activity (GO:0048018) | 2.31E-04 | 0.030279 | 2.08791 | 17.48102 |
| SH3 Domain Binding (GO:0017124) | 3.71E-04 | 0.03796 | 5.169185 | 40.82692 |
| Acrosin Binding (GO:0032190) | 4.35E-04 | 0.03796 | 40.5063 | 313.5576 |
| Calcium Channel Activity (GO:0005262) | 7.67E-04 | 0.057445 | 3.437791 | 24.65759 |
| Cytokine Activity (GO:0005125) | 0.001188 | 0.077782 | 2.466679 | 16.61529 |
| Chemokine Activity (GO:0008009) | 0.00193 | 0.09802 | 4.411399 | 27.57295 |
| Voltage-gated Calcium Channel Activity (GO:0005245) | 0.002155 | 0.09802 | 5.076741 | 31.17174 |

**Supplementary Fig. 4: Acute UNC1999 treatment on A375 cells elicited cell surface immune-related responses in A375 cells.** Enrichment of Gene Ontology terms for molecular processes for the set of genes downregulated genes after acute treatment with UNC1999 in (a) Parental cells and (b) Bottom-10 cells.

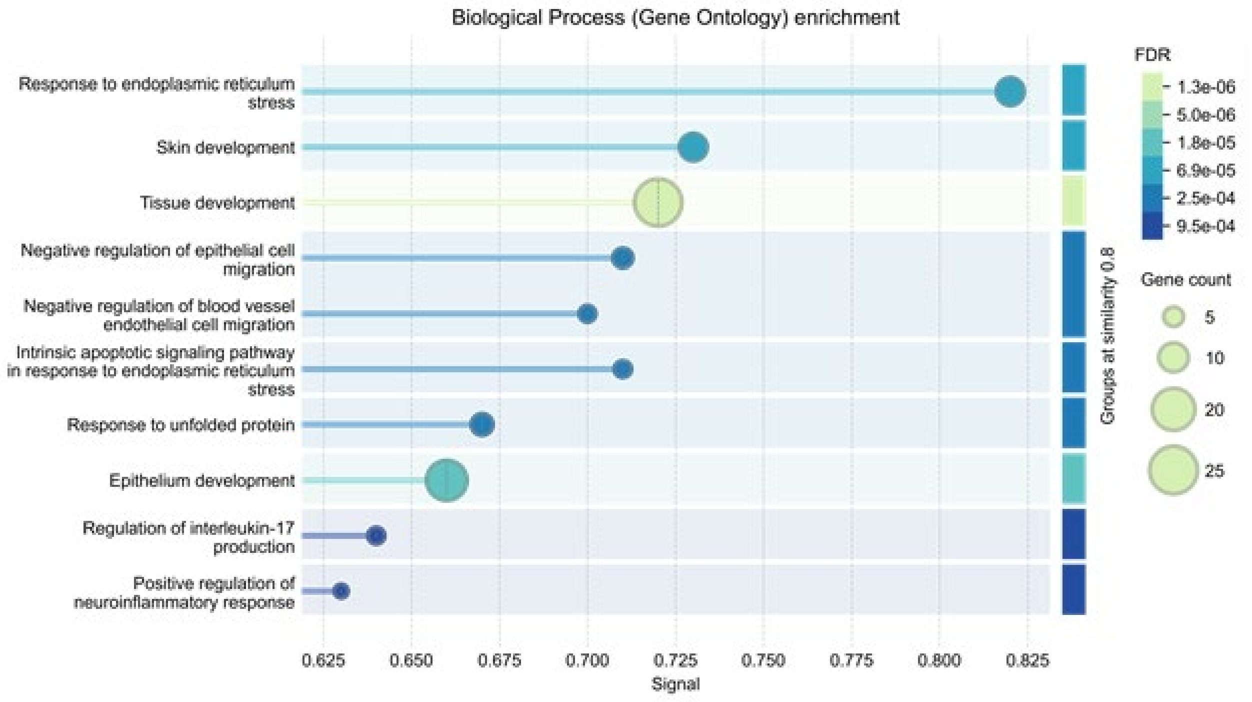


**Supplementary Fig. 5: Chronic EZH1 & 2 Inhibition during sequential migration leads to downregulation of genes involved in integumentary development**. GO terms (identified by StringDB) enriched among downregulated genes in UNC1999-treated Bottom-8 cells are involved in processes such as Skin development, tissue development, and epithelium development.
